# Push-and-pull protein dynamics leads to log-normal synaptic sizes and probabilistic multi-spine plasticity

**DOI:** 10.64898/2026.01.29.702571

**Authors:** J. Petkovic, M. F. Eggl, D. Pathirana, T. E. Chater, J. Hasenauer, S.O. Rizzoli, T. Tchumatchenko

**Affiliations:** Universitaet Bonn, Universitätsklinikum Bonn, Institute for Experimental Epileptology and Cognition Research; Institute of Neuroscience, CSIC-UMH; Life and Medical Sciences (LIMES) Institute & Bonn Center for Mathematical Life Sciences, University of Bonn; Laboratory for Synaptic Plasticity and Connectivity, RIKEN Centre for Brain Science, Saitama, Japan; Department of Physiology, Keio University School of Medicine, Tokyo, Japan; Department for Neuro- and Sensory Physiology, University Medical Center Göttingen and Center for Biostructural Imaging of Neurodegeneration, Humboldtallee 23, 37073, Göttingen, Germany

## Abstract

A typical neuron receives thousands of inputs and is able to adapt the strength of its synapses to store new information and meet ongoing computational demands. The synaptic response to plasticity induction is stochastic and spatially structured but is traditionally described by deterministic models representing the “average” dynamics. Growing experimental evidence indicates that not only the stimulation protocol determines the plasticity outcome but that the initial synaptic sizes, their fluctuations, and the spatial competition for the plasticity-relevant proteins play a decisive role. This probabilistic perspective makes it hard to predict the fate of a given synapse and requires a conceptual shift from a single synapse view to a probabilistic multi-spine competitive process where the plasticity needs and the available resources are considered together. Here, we propose a data-driven modeling framework able to predict collective plasticity outcomes along a dendrite based on the initial size, the number, and the spatial distance between simultaneously stimulated synapses. Our data analysis reveals a log-normal distribution of protein numbers for many plasticity-mediating proteins and shows that this log-normal protein allocation constrains and controls the collective plasticity outcome across multiple stimulated and non-stimulated synapses while preserving a global size distribution. Our findings highlight how local stochastic processes and global protein allocation rules give rise to synaptic plasticity outcomes, offering a new framework to understand and predict dendritic computation.

## Introduction

Synaptic plasticity is traditionally seen as a stereotypical and deterministic phenomenon where a synaptic change Δ*w*, to be added or subtracted to the initial synaptic strength *w*, is determined by the activity history, the cell type under investigation, and the intra- and extracellular environment of a synapse. Similarly, in artificial neural networks, training is based on the assumption that Δ*w* is determined by the global or local gradients but is independent of the weight *w* itself, and any weight fluctuations or internal state-dependence of the synaptic plasticity outcome are often neglected. However, experimental outcomes speak against the predictability and the deterministic nature of plasticity outcomes, pointing to a significant size variability in plasticity outcomes in response to the same protocol [1–3] and highlighting the probabilistic structure of synaptic sizes and their log-normal-like distribution [4, 5]. Furthermore, recent work suggests an important role for the initial (basal) state of the individual synapse as a driver of variability in plasticity outcomes [6, 7]. Recently recorded datasets with excellent spatial resolution at the protein and spine level [3, 8–11] put us now in the position to reconsider fundamental aspects of synaptic plasticity that were previously considered challenging.

Here, we build a data-driven model linking log-normal synaptic size variability and protein composition at the individual synapses to the variability in plasticity outcomes, identifying the molecular drivers underlying the varying plasticity outcomes at the level of individual synapses subject to the same induction protocol. Notably, our model can reconcile several previous seemingly contradictory results emerging from the dose-dependent application of FK506, which, interacting with both calcineurin and CaMKII*α*, can paradoxically increase [9] or reduce [12] the peak synaptic potentiation upon plasticity induction. Our new modeling framework can predict both an increase and a reduction of LTP as a function of the FK506 dose, as well as the balance between the initial synaptic protein composition and the phosphorylated protein fraction contributed by the induction protocol. Surprisingly, our results, moreover, predict that plasticity outcomes naturally comprise three groups of synaptic responses: synapses responding in the direction of the stimulation protocol, a non-responder fraction and a small synaptic fraction moving in the opposite direction. All three fractions have been previously reported in experiments [13, 14], but the last two fractions have been considered to be a failure in experimental execution, and were not considered to be a part of the natural plasticity outcome. Here, we show how all three fractions can co-occur and show that their relative probability is mediated by the initial molecular composition at the synapse and the stimulation protocol.

## Results

Dendritic spines are the location of the post-synaptic part of most excitatry synapses in the mammalian brain. These small femto-liter sized structures can change their volume, depending on the activity history of the synapse in question. Moreover, many synaptic measures positively correlate with spine volume, including PSD area [15], amount of both AMPA [16] and NMDA receptors [17], as well as the number of presynaptic vesicles [18]. Following synaptic stimulation, spines can show different plasticity responses, growing, shrinking, or exhibiting no change in their size [19] (Fig. 1). These responses are dictated by a variety of factors, related to the stimulus (e.g., its intensity, duration, or the number of stimulated spines) [10, 20], to the properties of each individual spine (e.g., their distance from the stimulus or its basal size) [1–3], and by the dendritic system as a whole [9, 21]. Importantly, these responses show a large degree of stochastic variability, with a significant inverse correlation between the synaptic tendency to potentiate and initial size or weight [14, 20, 22, 23]. To account for this diverse phenomenology, a multi-spine plasticity model needs to include specific functional components. First, it must account for the stimulus features, such as its intensity and the number of stimulated spines. Second, it must contain information about the spine locations (both homo- and heterosynaptic), and specifically about their distance from the stimulus locations. Third, it must include the temporal dependence of plasticity-induced changes. Fourth, it has to provide a sufficient degree of stochasticity to account for the inter-experimental difference in synaptic initial conditions.

**Figure 1.**
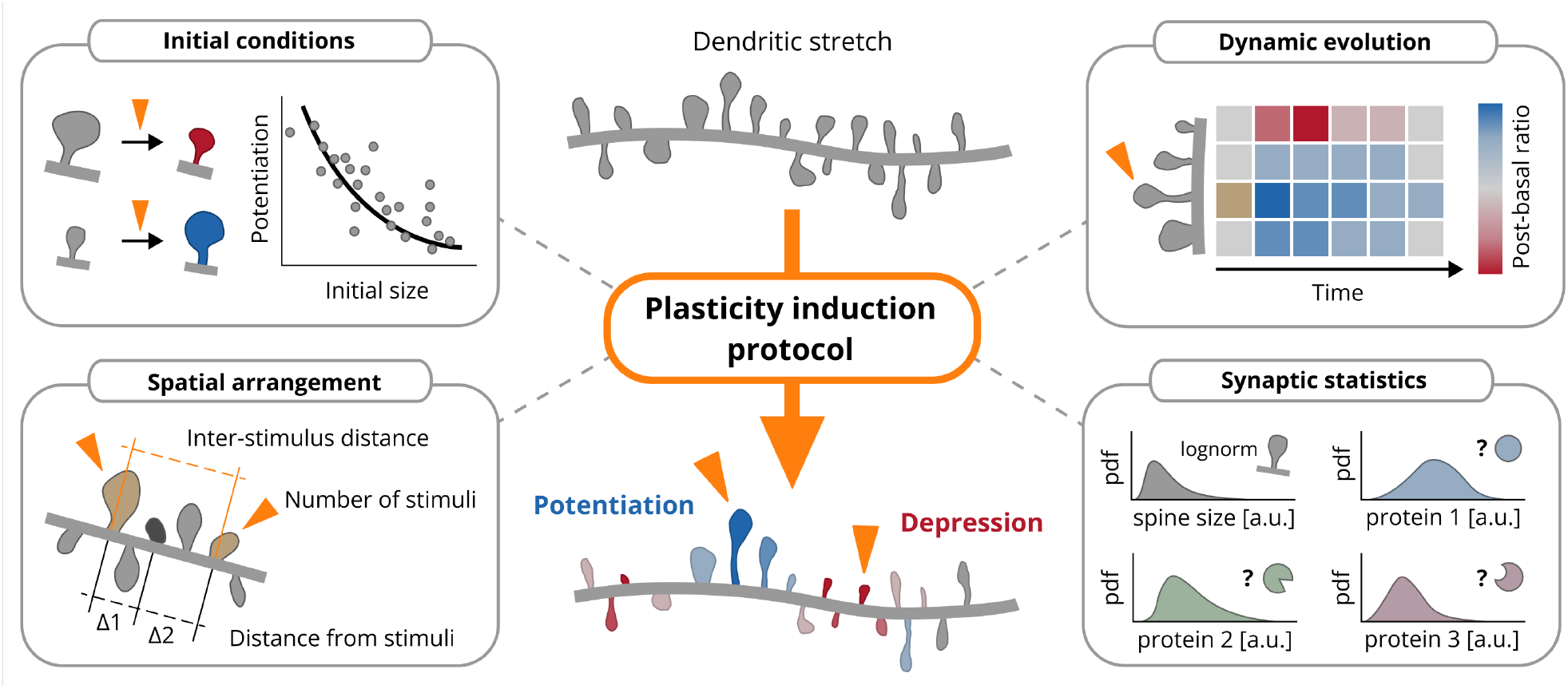
Overview of the parameters determining the multi-spine plasticity outcome. Top, left: initial synaptic size distribution. Bottom, left: spatial arrangement of the stimulation sites and parameters of the plasticity protocol. Top, right: elapsed time from the application of the stimulation protocol. Bottom, right: statistical properties of the proteins driving synaptic plasticity.

To build such a model, we start by considering a general synapto-dendritic system, focusing on the protein dynamics that take place throughout its domain (Fig. 2.c,b). Several different molecular processes have been shown to underpin synaptic size change during plasticity, giving rise to dynamics that occur on a range of different spatial and temporal scales [24–27]. Among these, phosphorylation-dephosphorylation cycles are specifically considered to control size changes on the minute-to-hour timescale [12, 13, 28–30], and has been successfully used by previous works to gain computational insight into various types of plasticity responses [31–33].

**Figure 2.**
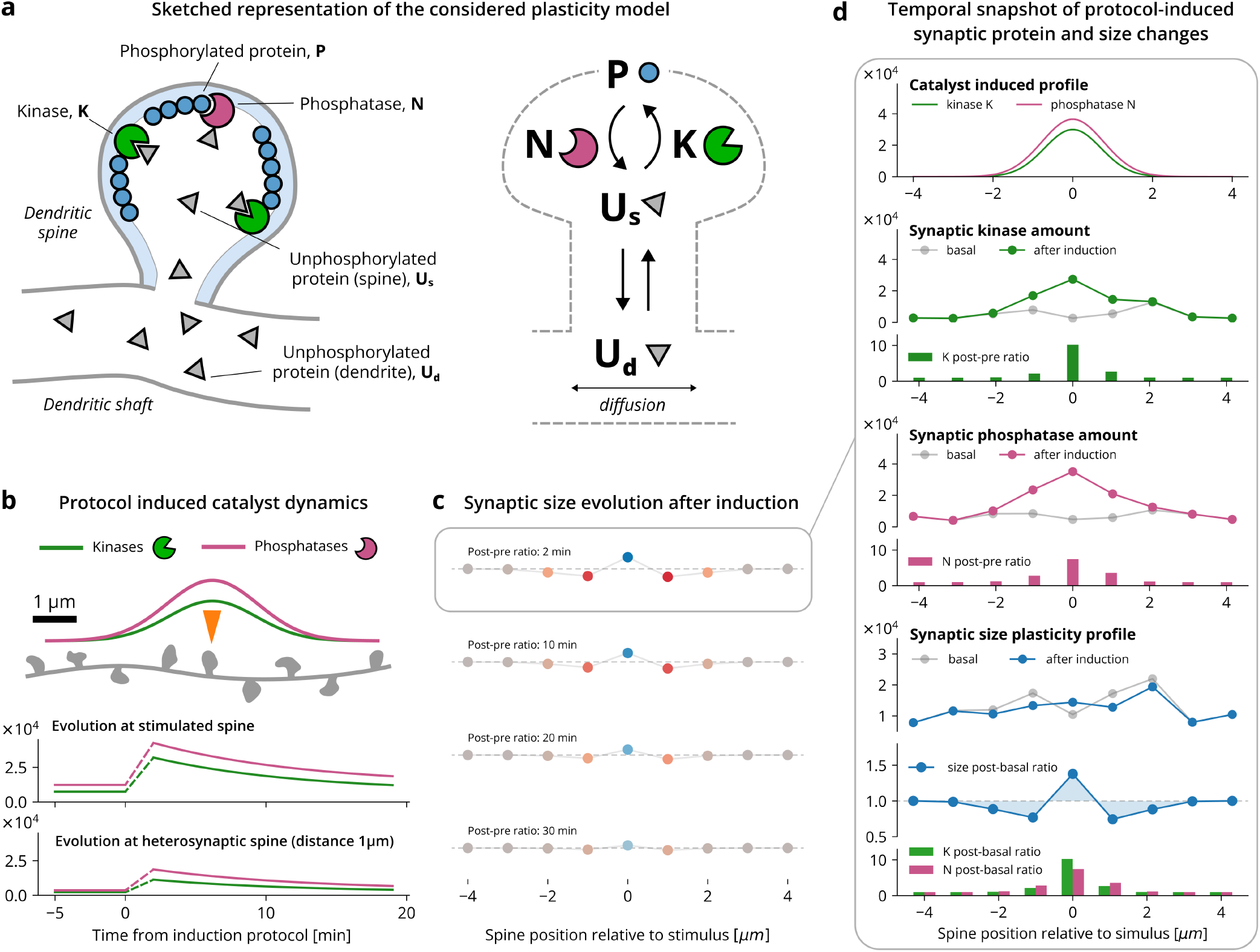
Overview of the molecular parameters used in our model and their role for plasticity outcomes. **a** Illustration of the model design and the push-pull dynamics of kinases (green) and phosphatases (pink) in the activation (phosphorylation) of the protein resources (blue circles) and their inactivated form (gray circles). The unphosphorylated protein pool *U* (gray) diffuses through the dendrite and enters the spines. Here it is reversibly converted to its phosphorylated counterpart *P* ^(*i*)^ by synaptic catalysts *K*^(*i*)^ and could be later deactivated by phosphatases *N* ^(*i*)^. **b** Stimulus induced synaptic catalysts. Each spine contains a unique, basal, stable amount of active kinases and phosphatases. This amount is increased by the plasticity protocol in a distance-dependent fashion (Gaussian kernel). After induction, the active catalysts decay exponentially to the original baseline. **c** Spatio-temporal profile of protocol-induced plasticity emerging from the proposed model. The protocol generates a Mexican hat plasticity profile, decaying exponentially in time and ultimately returning to baseline. **d** Temporal snapshot of the protocol-induced synaptic changes. Despite the amount of both active kinases and phosphatases being increased by the action of the stimulation protocol, the resulting plasticity profile allows both potentiation and depression as it is driven by the ratio of *K*^(*i*)^ and *N* ^(*i*)^.

We focus, therefore, on an abstract unphosphorylated synaptic protein resource, referred to as *U* . This resource is able to diffuse throughout the dendrite (dendritic fraction *U*_*d*_) and enter and fill the dendritic spines, which we index with *i* (synaptic fraction 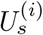). Within the dendritic spine, two general families of active catalysts are present: kinases (referred to as *K*^(*i*)^) and phosphatases (referred to with *N* ^(*i*)^). These two families regulate the interconversion between the unphosphorylated resource 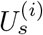 and its phosphorylated counterpart, denominated *P* ^(*i*)^, which is integrated into the synaptic structure and leads to the experimentally observed change in synaptic size [34, 35]. In our model, we assume that this phosphorylated resource is predominantly located inside the spines, and that a potential minor leakage into the dendrite can be neglected. This process, orchestrating the dynamics of *n* spines located on a dendritic stretch of length *L*, can be expressed as a system of differential equations:

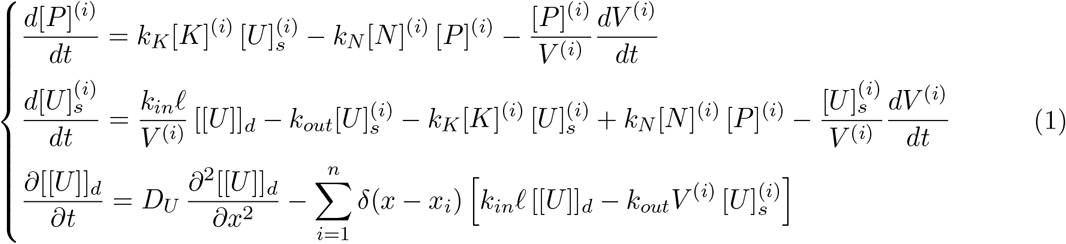

where *k*_*K*_ and *k*_*N*_ are the phosphorylation and dephosphorylation rates, *k*_*in*_ and *k*_*out*_ are the synapse-dendrite resource exchange rates, *V* ^(*i*)^ is the volume of the *i*-th spine, ℓ is a dimensional unit of dendritic length, and [[·]] and [·] are linear and volumetric density, respectively. It is crucial to notice the appearance of the *dV* ^(*i*)^*/dt* terms, accounting for the fact that synaptic sizes are not constant, impacting the concentration dynamics of the considered species.

This system, describing the introduced protein concentrations, allows us to derive a strikingly intuitive closed-form expression for the synaptic sizes *V* ^(*i*)^. The derivation (explained in the Methods and reported in mathematical detail in the Supplementary Information), relies on two observations reported extensively in the experimental literature (e.g., [36, 37]). First, that the concentration dynamics of the catalysts *K*^(*i*)^ and *N* ^(*i*)^ change slowly, compared to the timescales of the diffusion and phosphorylation processes. Second, that there exists a linear dependence between synaptic volume and phosphorylated protein content (represented in our model by *P* ^(*i*)^) [8, 38, 39]. Under these conditions, we carry out quasi-steady-state approximation and obtain the final expression for the synaptic size dynamics

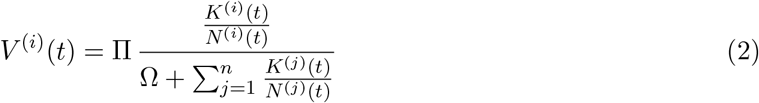

where Π is a constant proportional to the total amount of resources present in the dendrite and the spines, and 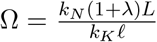 is a parametric constant accounting for dendritic length *L*, catalytic rates, and synaptic linear density *λ*. Notice that this equation does not depend on the concentrations of the catalysts *K*^(*i*)^ and *N* ^(*i*)^ but on their absolute values.

Some important functional properties can already be recognized in (2). First, different contents of active kinases and phosphatases directly translate to different spine sizes. Second, changes in the amounts of synaptic kinases and phosphatases induce a *local* change of synaptic size, while a change in the total amount of resources Π translates to a *global*, multiplicative change of all synaptic sizes. Third, since (2) is a homographic function of the ratio *K*^(*i*)^*/N* ^(*i*)^, synaptic sizes are structurally prevented from taking infinite value, as *V* ^(*i*)^ can at most reach the value of Π, when the ratio *K*^(*i*)^*/N* ^(*i*)^ tends to infinity. From a biochemical standpoint, this corresponds to a scenario where all the available resources have been sequestered by the *i*-th spine, completely depriving the rest of the synapto-dendritic system.

In this framework, the spatio-temporal effect of a plasticity protocol is mediated by the changes in active *K*^(*i*)^ and *N* ^(*i*)^, which, evolving in time, drive the observed changes in synaptic size. As last step, therefore, we model the time evolution of the catalysts *K*^(*i*)^ and *N* ^(*i*)^ in relation to a stimulation protocol. In accordance with the literature [9, 31, 36, 40, 41], we assume that (1) immediately after induction, the number of activated catalysts spikes to a new, higher amount, (2) that it then decays exponentially with time to its basal value, and (3) that the catalyst activation effect depends on the distance from all the stimulations in a Gaussian-shaped fashion (Fig. 2.e). The equations describing these dynamics are

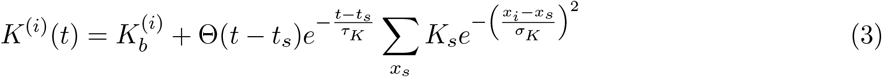

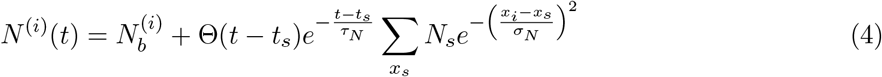

where *t*_*s*_ and *x*_*s*_ are the time and synaptic locations of plasticity induction, Θ is the Heaviside theta function, *τ*_*K*_ and *τ*_*N*_ are the catalyst decay timescales, 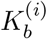 and 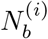are the basal active catalyst values at each spine *i, K*_*s*_ and *N*_*s*_ are the homosynaptically induced catalyst amounts, and *σ*_*K*_ and *σ*_*N*_ are the spatial induction decay scales.

A graphical representation of the model behaviour is shown in Fig. 2.c-d, where plasticity induction is carried out stimulating the central spine of a small dendritic stretch. We can see that both homosynaptic potentiation and heterosynaptic depression can emerge in a Mexican-hat like fashion [3, 19], due to the interplay between the stochastic, basal synaptic catalyst amounts and the deterministic distance-dependent induction provided by the stimulus (Fig. 2.g). Ultimately, progressing with time, the induced changes vanish, and the spines return to their initial sizes.

To be able to use this framework and investigate the mathematical principles driving synaptic plasticity, we must first infer its optimal parametrization. To this end, we fit our model to the experimentally observed synaptic size dynamics reported in [9], together with additional data obtained by repeating this same experiment with 5 closely located stimulations (details reported in the Optimization section of the Methods). In the next sections, we will start by analyzing the results of this fit, and then proceed to use our framework to simulate previously reported plasticity experiments, disentangling the functional principles leading to the diversity of their outcomes.

### Synaptic size log-normality can emerge from synaptic trapping and diffusive dynamics

The first question we are interested in addressing with our model regards the basal statistical properties of the synaptic plasticity-mediating proteins. As shown by our equations, the variability of these proteins directly impacts the ability of each spine to compete for resources, shaping both basal synaptic sizes and their response to plasticity induction. Ultimately, this mechanism leads to the emergence of the population statistics of synaptic sizes, which have been extensively characterized from an experimental point of view but still represent an open challenge for mechanistic modeling.

We start by focusing on the inferred values of basal synaptic catalysts 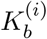 and 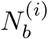. Log-normal pat-terns have been observed in relation to a number of synaptic metrics [4], suggesting that log-normality can arise from a fundamental process driving protein dynamics inside dendrites. Importantly, several recent studies [11, 42] also show that log-normality can be uncoupled from neuronal activity, arguing that its emergence could arise not from a top-down information encoding feature but from a basic intracellular process.

Following this line of reasoning, we test the fitted distributions of 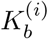 and 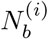 and find that, in-deed, they do not show statistically significant difference from a log-normal distribution (Fig. 3.a,b). Moreover, the values of these two catalyst families show a substantial degree of correlation across spines (Fig. 3.k), with a highly significant Pearson *r* value of 0.84. These two findings (log-normality and correlation between 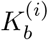 and 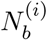) suggest that a good candidate distribution for these two cat-alyst families is a joint, bivariate log-normal distribution, of which each dendritic spine represents an independent realisation.

**Figure 3.**
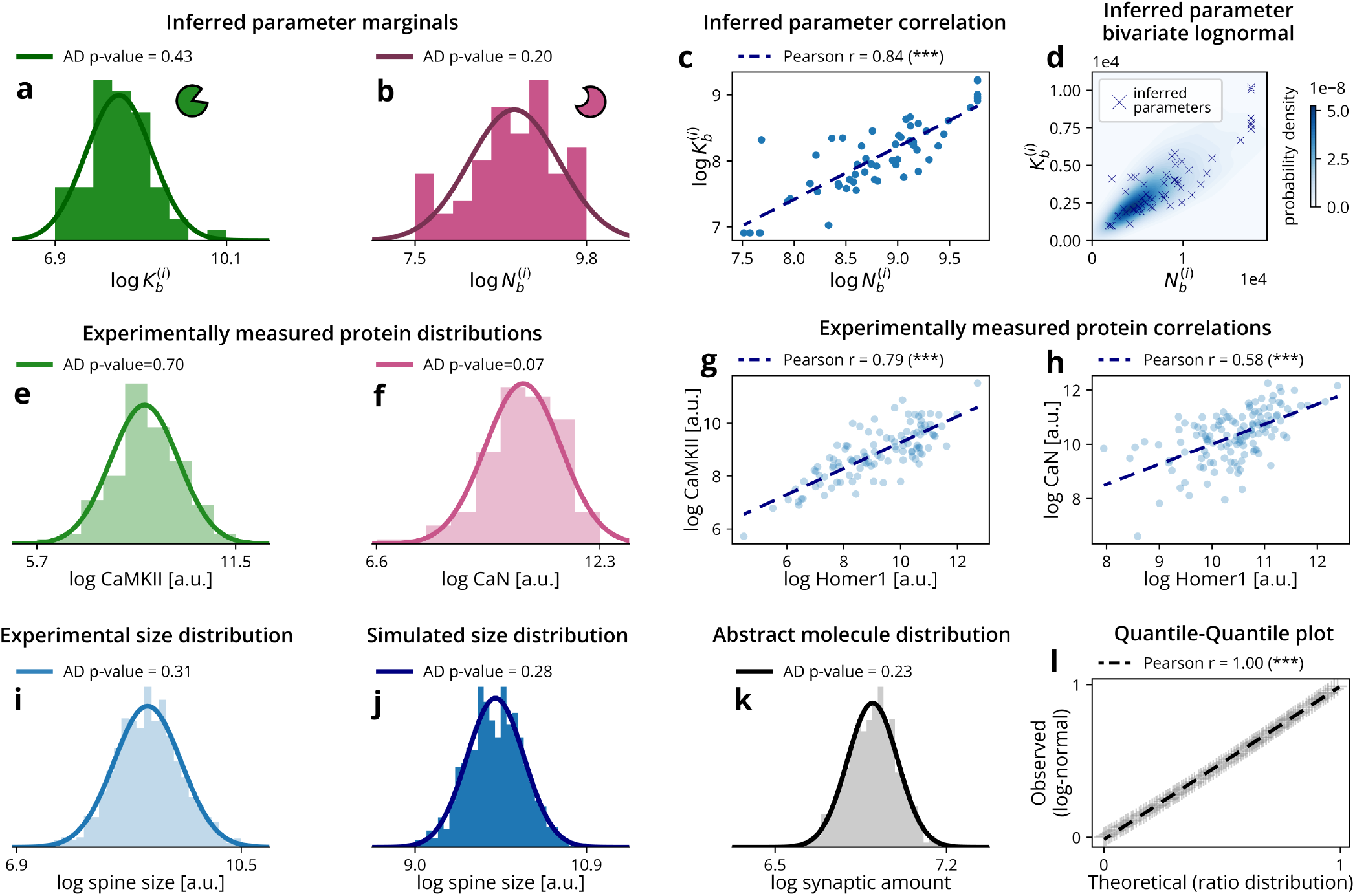
Statistical features of the synaptic plasticity-related proteins. When presented, log-normality is assessed by testing the logarithms of the data for normality with the Anderson-Darling (AD) test. A resulting p-value higher than 0.05 denotes compatibility with a log-normal distribution. **a**,**b** The inferred optimal values of synaptic 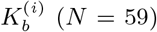 and 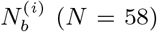 each follow a log-normal compatible distribution. **c** The logarithms of the inferred synaptic catalyst amounts show a strong, significant linear correlation. **d** Graphical representation of the bivariate log-normal describing the distribution of the estimates of 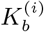 and 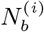 **e**,**f** Experimentally observed distributions of synaptic Ca2+/calmodulin-dependent protein kinase II (CaMKII, *N* = 110) and calcineurin (CaN, *N* = 133) across spines. **g**,**h** Correlation between synaptic Homer1 and synaptic CaMKII/CaN signals respectively. **i**,**j** Experimentally observed (*N* = 1105) and model simulated (*N* = 1000) distributions of synaptic sizes. Both results are compatible with a log-normal distribution. **k** Synaptic steady-state distribution of a freely diffusing abstract molecule, showing strong compatibility with a log-normal distribution (*N* = 1000). **l** Quantile-quantile plot (QQ plot) comparing the known theoretical distribution of the diffusive molecule (ratio distribution) with a fitted log-normal. From both the visual inspection and the linear correlation coefficient, the two distributions are indistinguishable.

To confirm the robustness of the log-normality across different model-fits and experimental data we conducted several cross-checks.

First, we confirmed that a substantial fraction of optimization runs were converging to the best local minimum (Fig. S2) and proposed synaptic estimates for *K*^(*i*)^ and 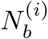, all showing log-normal distributions. Moreover, focusing on each spine, we also observed that the emerging distributions across runs were compatible with a log-normal for the large majority of the spines (Fig. S6). These facts strongly suggest that the parameter combinations in our model that recover the experimentally observed synaptic plasticity dynamics are consistent with a log-normal distribution of catalytic proteins across spines.

To corroborate the validity of this observation, we investigated whether the synaptically localized proteins also show a distribution compatible with a log-normal distribution. To this end, we analysed two experimental datasets, presented in [8] and [43] super-resolution protein quantification is conducted in both pre- and post-synaptic terminals. Encouragingly, we found that numerous different plasticity-related proteins showed a log-normal distribution across both boutons (Fig. S13) and dendritic spines (Fig. S12), in this latter case with a high degree of correlation with the scaffolding protein Homer1. The log-normal statistics were also present in the two model-relevant synaptic catalysts, Ca2+/calmodulin-dependent protein kinase II (CaMKII) and calcineurin (CaN) (Fig. 3.e-h), which can be considered two major biological counterparts of the model’s kinase and phosphatase families *K*^(*i*)^ and *N* ^(*i*)^. Moreover, their correlation with Homer1 allows us to derive theoretical bounds for the correlation between the inferred 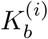 and 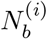, the result of which is compatible with the value obtained in the optimization (Pearson *r* = 0.79, theoretical bounds [0.05, 0.92], computed following the equation in [44]). Taken together, these observations support the hypothesis that the model’s catalytic families follow a bivariate log-normal distribution.

As final cross-check for lognormality, we set out to find a candidate mechanism that can lead to the emergence of this log-normal distribution across spines for a general neuronal protein. To this end, we use again the stochastic simulation of a linear dendrite, where a molecule species diffuses throughout the dendritic shaft, exiting and entering dendritic spines (more details are provided in the relevant section of the Supplementary Information). Under very general conditions, we find that the resulting synaptic distribution of this abstract protein resource is indistinguishable from a log-normal distribution (Fig. 3.k-l), and that this log-normality emerges from the basic diffusion-and-exchange process in which a synapse-to-synape variability in synapse-to-dendrite exchange rates is present.

Having found a biologically plausible mechanism and its theoretical description, we conclude that the catalytic distributions of 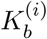 and 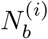 follow a bivariate log-normal distribution. This has an immediate modelling consequence. Given the log-normality of the marginals 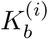 and 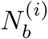, the ratio 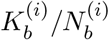 also follows a log-normal distribution, directly leading through the model equation (2), to log-normality of synaptic sizes (Fig. 3.i,j, derivation in the relevant section of the Methods).

In summary, after optimizing our model (2) to fit our plasticity data, we find that the basal catalytic synaptic distributions 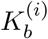 and 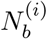 follow a bivariate log-normal distribution. To verify the robustness of this result, we verify the presence of log-normality in experimentally observed synaptic protein distributions and catalysts, and show that, under general assumptions, the universal process of diffusion-exchange leads to the emergence of such log-normality. Finally, we show that log-normality of synaptic proteins directly translates to the widely observed log-normality of synaptic sizes as a ratio of log-normal distributed variables.

### How synaptic size shapes the plasticity response

After characterizing the distributional properties of the basal synaptic catalytic families, we set out to understand how these properties impact synaptic size change when a stimulation protocol is applied, starting from the perspective of a single, stimulated spine.

Synaptic response to plasticity induction is characterized by a high degree of variability, which stems from biological, methodological, and observational noise [2, 22, 45]. Several experimental and computational works have reported an inverse power-law-like relationship between the magnitude of synaptic potentiation and its initial size [2, 7, 20, 22], but the origin of this dependence has never been conclusively identified.

We address these questions by simulating an experiment where plasticity is triggered at a single spine on a dendritic stretch, using a potentiating protocol in line with [9] (this choice comes from our parametrization being inferred on this specific dataset). Starting from different synaptic initial conditions (different 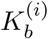, 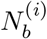, and, therefore, synaptic sizes), we see a substantial degree of response variability, with different repetitions of the same experimental protocol leading to different plasticity behaviours (Fig. 4.a). Importantly, while, on average, the stimulated spine shows statistically significant potentiation, it is not impossible to observe depression as well, despite the protocol in [9] being potentiating (Fig. 4.a bottom and middle row).

**Figure 4.**
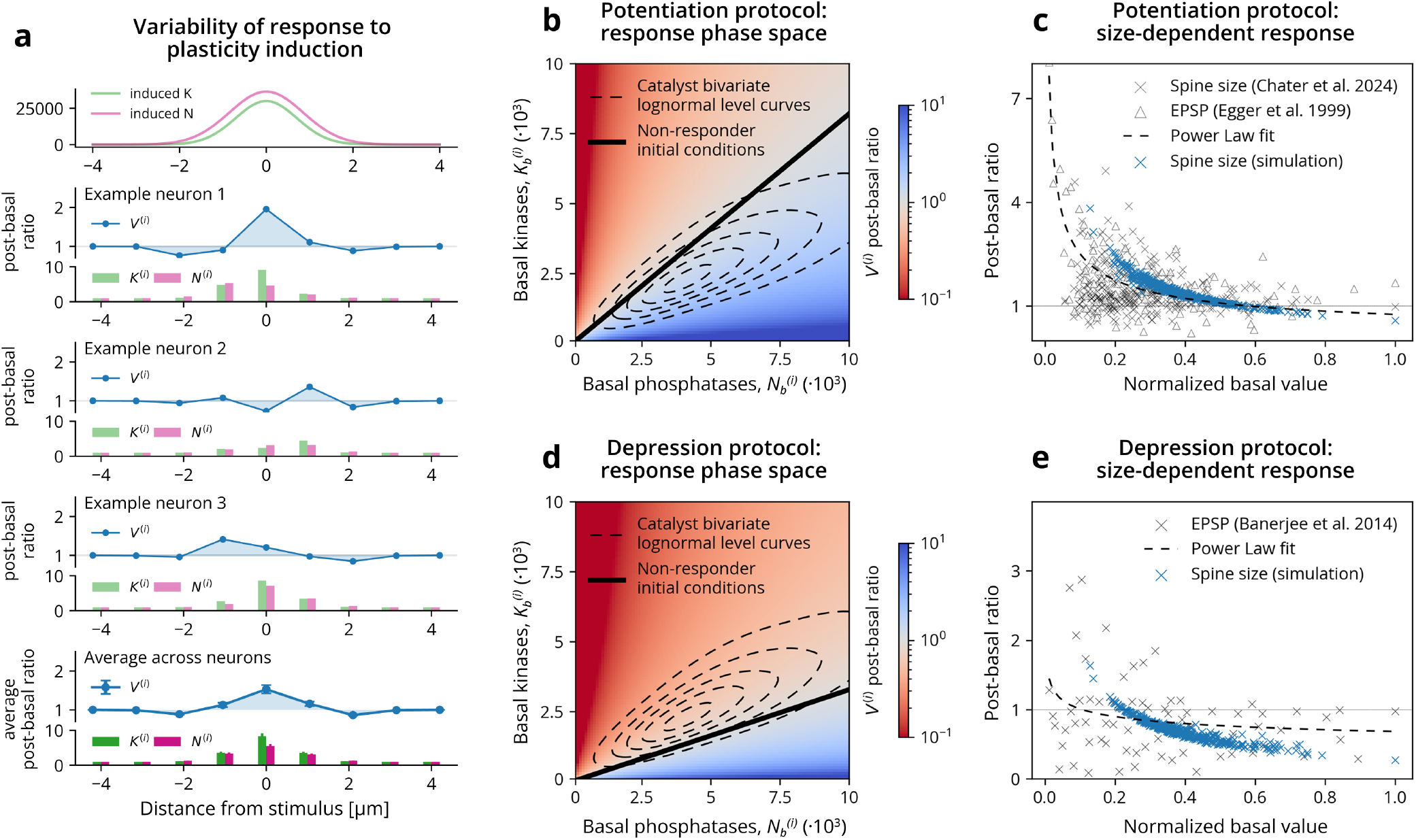
Plasticity outcome dependence on basal synaptic statistics. **a** The application of a synaptic potentiation protocol provides a fixed contribution of active synaptic kinases and phosphatases (upper panel). The synaptic response, however, is determined by the sum of this deterministic quota with the stochastic basal amount, and varies significantly across experiment repetitions (second, third,and fourth panel). On average, however, a well-defined spatial plasticity profile emerges, uniquely determined by the stimulus type and the synaptic protein statistics. **b** For each basal synaptic condition (a pair 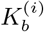 and 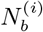), the model enables calculation of the average synaptic response (*P* ^(*i*)^post-basal ratio) to a given stimulus protocol. This response depends on the synaptic catalytic distribution, with a clear distinction between potentiating (blue) and depressing (red) regions. Between these regions, a set of initial catalytic values leads to no change after stimulus (black line). **c** Several synaptic measures (spine size, EPSP) show an overlapping trend between their relative variation (post-pre ratio) and their normalized basal value. Our simulations also lead to a very similar behaviour, closely following a power-law fit to data proposed in [7]. **d** same as panel b with a depressing protocol. Notice that the majority of the spines fall in the red region, allowing the protocol to be considered depressing in an experimental setting. **e** Same as panel c with a depressing protocol. The simulation does not obey the power-law fit to data, but qualitatively recovers the inverse relationship between spine size and potentiation/depression, allowing the potentiation of small spines.

This variability in responses directly derives from the stochastic nature of the basal 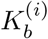 and 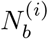 distributions, and can be clearly understood by resorting to a phase space representation (Fig. 4.b,c).

In this representation, a spine before stimulation is uniquely identified by a vector, a pair of values 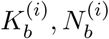, corresponding to its basal catalytic content. Importantly, this vector corresponds to a specific synaptic size. As discussed in the previous section, biological synapses do not uniformly fill this phase space, but follow a bivariate log-normal distribution (Fig. 3.d and dashed lines in Fig. 4.b,d). For a given stimulus, our model can predict the average maximal response for each point of the phase plane (red-blue color code in Fig. 4.b,d), showing that the same stimulus can lead to both potentiation and depression, as a function of the initial synaptic size. The transition between potentiation and depression is smooth, and traverses a line of synaptic sizes that do not show any response to the plasticity inducing protocol. We call these family of spines *non-responders* (Fig. 4.b and d, black lines). Since we have provided a closed-form solution for our model, we can also find the explicit equation of this line, as it represents the 0 level-set of average synaptic change (see Methods for derivation). From a practical standpoint, conducting multiple trials of a synaptic plasticity experiment consists precisely of fixing the stimulus parameters (the color background) and sampling the catalytic phase space from the bivariate catalytic distribution (dashed level curves); the average response (the color corresponding to the mean of the bivariate log-normal) is then what defines the protocol as potentiating or depressing.

The relationship between the stimulus features (the color background) and the synaptic distribution of catalysts (dashed level curves) following (2) is, ultimately, what leads to the emergence of the inverse relationship between synaptic potentiation and synaptic initial size. To validate this mechanism, we again turn our attention to experimental observations, describing synaptic size [9] and synaptic weight evolution [22] after a potentiating protocol is applied (Fig. 4.c). After normalizing the basal values of each dataset (dividing them by their maximum), we see that the simulated variations of *V* ^(*i*)^ show good agreement with the experimentally observed plasticity changes. Crucially, we also see that the simulation closely follows the power-law fit traditionally proposed as a descriptive model [7], recovering the three types of qualitatively different responses (potentiation, depression, and lack of change) with one single mechanism.

So far, we have reasoned that the difference between plasticity protocols resides only in the value of the stimulus-related parameters. This implies that our model should be able to reproduce the synaptic response to an LTD protocol after a suitable reparametrization (e.g., a reduction of the *K*_*s*_ or increment of the *N*_*s*_ values) (4.d). We test this prediction against experimental data taken from an LTD induction experiment [2], and report the outcome in Fig. 4.e. Despite showing less adherence to the experimental data, the model still correctly predicts the existence of the inverse relationship between synaptic change and initial synaptic size, associating overall depression with a crucial fraction of very small synapses undergoing potentiation.

In summary, we show that resource competition through phosphorylation can explain the elusive variability that characterizes synaptic responses to plasticity induction. Moreover, this mechanism is able to predict the experimentally observed inverse relationship between synaptic size and tendency to potentiate, ultimately leading to the same induction protocol to be able to result in all three possible changes in synaptic size (growth, shrinkage, and no change).

### A single mechanism for a multitude of plasticity profiles

In the previous section we have investigated the impact of the basal catalytic distributional properties on the response that a single spine shows to a plasticity-inducing protocol. We now ask ourselves how these properties impact synaptic plasticity from a more general, multi-spine perspective.

We start by testing our model, and simulate the synaptic size dynamics generated by inducing plasticity simultaneously at a 7 spine cluster spanning 16 *µm* . This configuration follows one of the experimental protocols reported in [9] that that was not used for our optimization. The simulation matches very closely the dynamics observed in the experiment (Fig. 5.a), with synaptic change being particularly evident at 2 minutes after the induction. Three qualitatively different behaviours emerge as a function of the distance from the closest stimulation (Fig. 5.a, middle panel). The stimulated, as well as the spines very close to the stimulation (∼ 2 *µm*) undergo potentiation, spines located at an intermediate distance (∼ 2 − 4 *µm*) undergo depression, and, lastly, spines that are further than 4 *µm* do not show any significant change in size. These effects decay over time, with synaptic sizes returning to baseline at the final timepoint of 40 *min* post-stimulation (Fig. 5.a, right panel). In addition to the average spatiotemporal plasticity dynamics, the model is also able to match the experimentally observed right-skewed distribution of post-basal size ratios at 2 minutes after the induction (Fig. 5.a, Kolmogorov-Smirnov test *p* = 0.13, *N* = 50), showing a small, albeit crucial, fraction of stimulated spines that undergo depression, in accordance with the mechanism described in the previous section.

**Figure 5.**
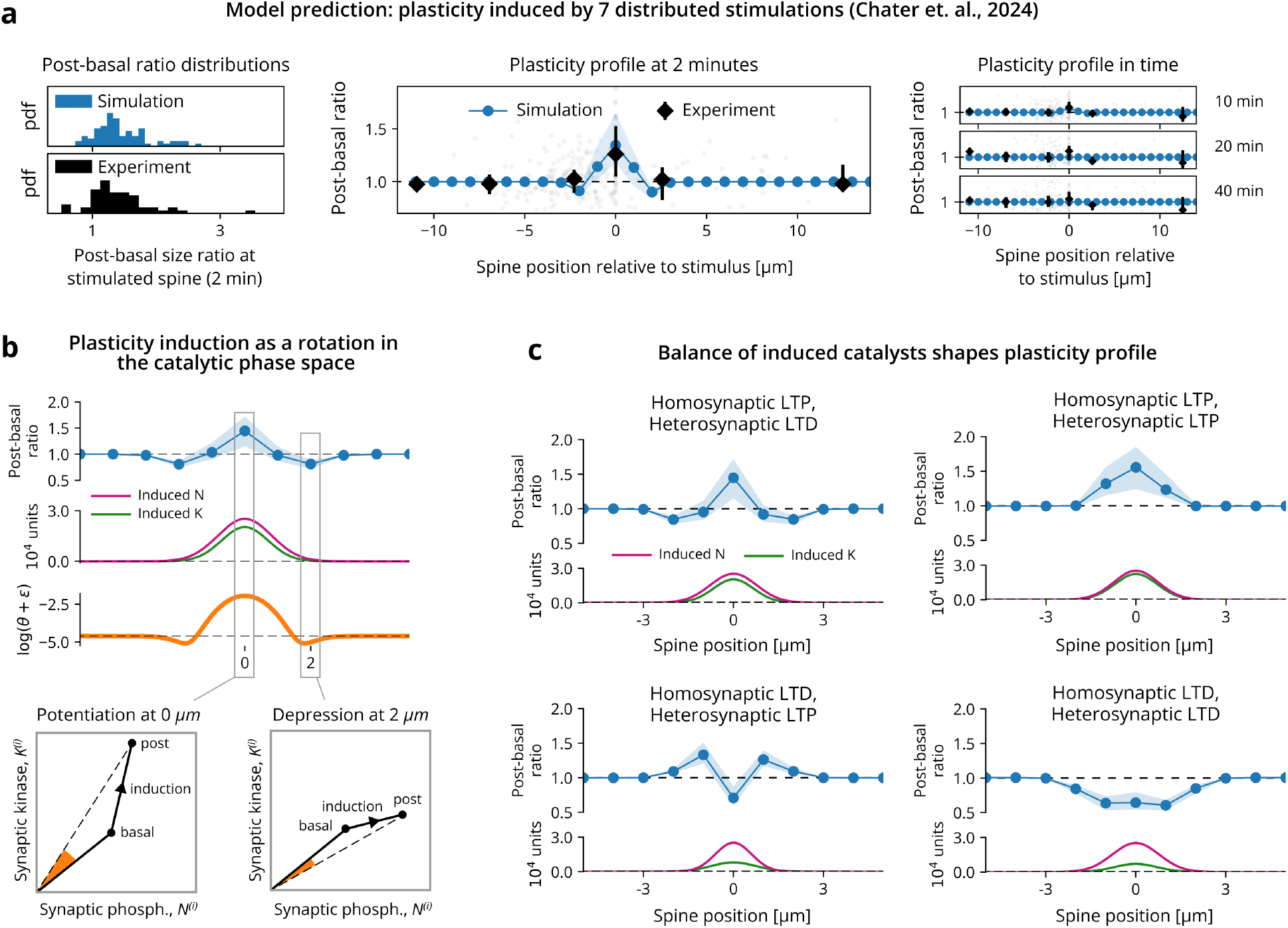
The role of the catalytic activation profile in inducing multi-spine plasticity. Where reported, experimentally observed and simulated synaptic size ratios are expressed in terms of median and interquartile ranges, as the underlying distributions are not Gaussian. **a** Middle and right panel: model prediction comparison with the test dataset from [9]. Spatial profile and temporal evolution of plasticity induced synaptic sizes closely match the experimentally observed dynamics. Left panel: our model is also able to correctly predict the skewed distribution of the stimulated post-basal size ratios, emerging 2 minutes after induction (Kolmogorov-Smirnov *p* = 0.13, *N* = 50). **b** Plasticity induction at the central spine leads to the emergence of a Mexican-hat response profile. Upper panel: synaptic size change profile, together with the catalytic activation profile and the consequent *θ* profile are shown in function of space. For better visualization, we plot the logarithm of *θ* offset by a small value *ϵ*, highlighting the presence of both positive and negative values (higher or lower than *ϵ*, in our plot). Lower panel: at different distance from the stimulus, spines get induced different amounts of kinases and phosphatases. The rotation angle *θ* between the “basal” synaptic conditions and the “post” synaptic conditions is proportional to the induced kinase/phosphatase ratio, and dictates the sign of the synaptic response (potentiation of clockwise rotations - *θ >* 0, and depression otherwise). **c** Gaussian catalytic activation profiles are able to recover the full spectrum of experimentally observed plasticity profiles [19]. Assuming constant phosphatase induction (height of pink line), the amount of potentiation is proportional to the induced kinase amount (height of green line), switching from a fullly potentiating profile (high kinase induction), to an intermediate Mexican-hat shaped profile (intermediate kinase induction), to a fully depressive profile (low kinase induction). The direction of the Mexican-hat induction profile depends, additionally, on the spread of the induced *K* and *N* (width of the respective Gaussians), with peripheral depression emerging from wider phosphatase induction, and peripheral potentiation from the opposite scenario.

The origin of these three, qualitatively different responses can be understood by resorting once again to the catalytic phase plane representation (Fig. 5.b). As illustrated in the previous section, on this phase plane each spine represents a vector of basal catalytic contents 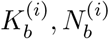 (“basal” point in Fig.5.b, lowermost panels). The stimulus acts as the addition of a new vector to these basal coordinates

(indicated as “induction” in the figure), with components depending on the distance between the considered spine and the stimulus locations (this spatial dependence is given by (3)). This addition, leading from the “basal” to the “post” coordinates, induces a rotation *θ* of the synaptic coordinates, which is, ultimately what describes the synaptic response to plasticity: clockwise rotations (negative *θ*) correspond to depression, counterclockwise rotations (positive *θ*) correspond to potentiation, and no rotation (*θ* = 0) indicate no change in synaptic strength. The origin of this principle becomes clear when considering that *θ* is a monotone function of the synaptic catalytic ratio *K*^(*i*)^*/N* ^(*i*)^ (*θ* = arctan(*K*^(*i*)^*/N* ^(*i*)^)), with positive or negative rotations corresponding to an increase or reduction of this ratio, and, consequently, of synaptic strength.

The rotation mechanism acts jointly on homo- and heterosynaptic spines (Fig. 5.b upper panel), with a Mexican-hat size plasticity profile deriving from an underlying Mexican-hat profile of *θ*, in turn originating from the Gaussian spatial profiles of catalytic activation induced by the stimulus. The response difference between homo- and heterosynaptic spines, therefore, can be described with a single mechanism, driven by different, distance-dependent values of induced catalysts.

Importantly, this same principle is not limited to the Mexican-hat plasticity profile, but can recover all four varieties of combined homo- and heterosynaptic plasticity outcomes reported in [19] (Fig. 5.c). Two factors, in particular, play a fundamental role. First, the intensity of the stimulus, proportional to the maximal amount of induced kinases *K*^(*i*)^ [37], is able to linearly switch the plasticity response from a fully depressive regime to a fully potentiating one, passing from an intermediate response showing homosynaptic potentiation associated with heterosynaptic depression. Second, the catalytic activation spread constants *σ*_*K*_ and *σ*_*N*_, dictated, e.g., by the compartmentalization features of the dendrite, are critical for shaping the direction of the Mexican hat profile, with higher values of *σ*_*N*_ leading to heterosynaptic depression, and higher values of *σ*_*K*_ to heterosynaptic potentiation.

Different synaptic plasticity responses arise not only from different stimulation protocols but are also associated with differences in the basal properties of the neurons undergoing plasticity [19]. Noticing that different neuronal properties entail differences in bas al catalytic distributions, we are able to show that this phenomenon can emerge directly from changes in these distributions. For a fixed stimulus, the plasticity response profile can show a spectrum of responses, from fully potentiating (low basal 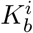or high basal 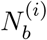) to fully depressing (high basal 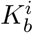 or low basal 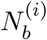, Fig. S14).

In summary, we show that homo- and heterosynaptic plasticity can be reconciled under a common multi-spine plasticity mechanism, grounded on resource competition through phosphorylation. Through the interplay between the basal distributions of synaptic catalysts and the space-dependent activation induced by the stimulus, this framework not only clarifies the link between stimulation and synaptic response, but also shows how qualitative differences in this response can emerge from quantitative differences in basal neuronal conditions.

### Phosphatase inhibition leads to bidirectional changes in potentiation

After understanding how resource competition and catalyst induction drive multi-spine plasticity, in our final section, we focus on how this mechanism is shaped by a molecular perturbation introduced during the experiment.

The effect of FK506 (tacrolimus), an inhibitor of the synaptic phosphatase calcineurin, has been extensively investigated throughout the literature in regard to its effects on synaptic plasticity, leading to conflicting observations. Previous work from our lab [9] has shown that CaN inhibition leads to an overall increase of induced potentiation, with higher and more lasting synaptic strengthening occurring both at stimulated and hetero-synaptic spines. This effect is intuitive, since a reduced amount of synaptic phosphatases should lead to an increase in calcium-induced protein phosphorylation and, consequently, overall synaptic potentiation. An opposing result, however, is presented in [12]. Here, application of FK506 leads to a reduction of maximal potentiation in a dose-dependent fashion. The authors explore several possible reasons that could explain this surprising effect, considering, among others, qualitative differences in the calcium signalling cascades induced by NMDAR and voltage gated calcium channels (VGCCs).

We propose a unified interpretation, able to reproduce these conflicting observations from a common dynamical framework (Fig. 6.a). The increase or the reduction in potentiation derive from an increase or a reduction of the ratio between basal and activated catalysts with respect to the control condition. Two factors drive this change. First, the addition of FK506 impacts the basal amounts of active CaMKII and calcineurin, increasing the first and reducing the second [37, 46]. Second, both FK506 and the difference in the calcium channels driving plasticity (NMDARs in [9] and VGCCs in [12]) modulate the stimulus-induced activation of calcineurin, due to inhibition and due to a quantitative difference in the emerging calcium dynamics. All these effects can be transparently implemented in the model, by changing the value of three parameters describing the corresponding catalytic features (Tab. S3). We start by reproducing the effect observed in [9], and modify the optimal model parameters to account for the effects of the addition of FK506, in accordance with previous experimental reports [37, 46, 47]. This strategy not only solves the technical challenge of fitting *de novo* a new optimal parameter set (impossible for such a small dataset), but also corroborates the claim that our model is grounded on an interpretable link between its parameters and their biochemical counterpart. Encouragingly, we find that the blocked model (Block POT) correctly reproduces the observed outcomes in [9], with higher and longer-lasting potentiation both at stimulated and surrounding spines (Fig. 6.c-f). We are also able to reproduce the slight increase in basal spine sizes observed after the FK506 has been applied to the culture (Fig. 6.c). In agreement with [37], this arises from an increase in average basally active kinases *µ*_*K*_ combined with a reduction of basally active phosphatases *µ*_*N*_ .

**Figure 6.**
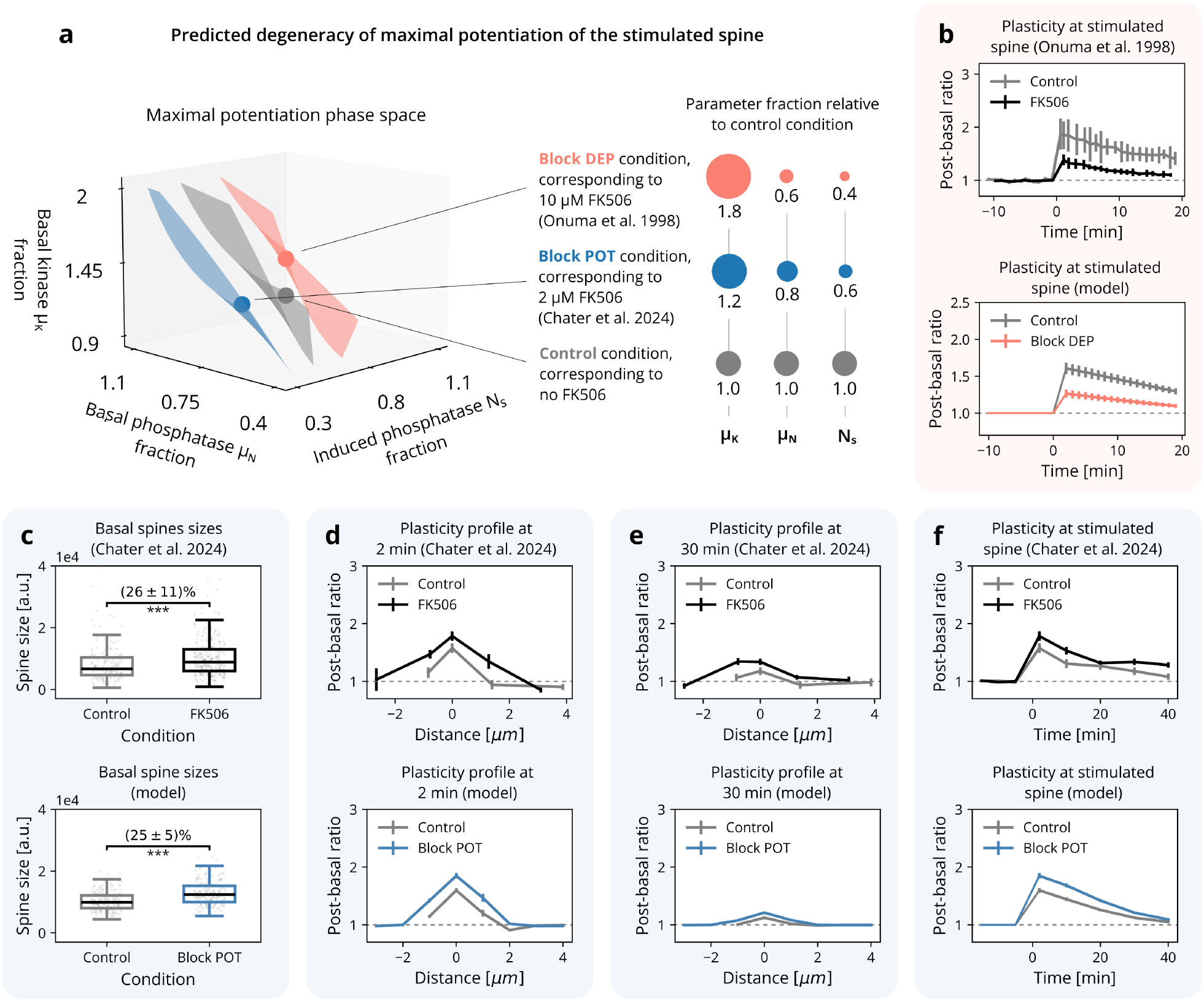
Differential FK506 inhibition of basal and induced catalysts leads to bidirectional effects on synaptic plasticity. Data is presented as mean *±* SEM, for compatibility with the considered experimental work. **a** Average maximum potentiation level surfaces (isosurfaces) in function of the average basal phosphatases *µ*_*N*_, average basal kinases *µ*_*K*_, and the induced phosphatase quota *N*_*s*_ (reported as fractions of the control parametrization). The surfaces correspond to parameter values that lead to the same average maximum potentiation. Different FK506 concentrations, as well as different types of biological conditions (e.g., different calcium channels), translate to distinct points in this parameter space. Importantly, due to the existence of these smooth isosurfaces, the model predicts that multiple different experimental conditions could lead to the same observations, potentially indistinguishable from the control condition (points on the gray surface). **b** Upper panel: experimental results reported [12]. The addition of FK506 reduces synaptic potentiation following plasticity induction. Lower panel: model reproduction, obtained with the parametrization Block DEP. **c-f** Upper panels: experimental results reported in [9]. The addition of FK506 induces a significant increase of the basal synaptic sizes (Kolmogorov-Smirnov *p*-value *<* 0.01, panel a), together with an of boost of synaptic potentiation after protocol induction, both temporally and spatially. Lower panels: model reproduction, obtained with the parametrization Block POT. Remarkably, due to the model’s explicit dependence on synaptic initial conditions, we are able to recover also the effect of FK506 on basal spine sizes (Kolmogorov-Smirnov *p*-value *<* 0.01).

After accounting for the difference in applied concentrations of FK506 (2 *µM* in [9], 10 *µM* in [12]), we can repeat the procedure described above, and obtain the results presented in [12]. As before, we are able to reproduce the correct plasticity behaviour, this time with the addition of FK506 inducing a reduction of potentiation at the stimulated spines (Fig. 6.b, Block DEP). In fact, our model indicates that potentiation change shows a continuous, albeit nonlinear, dependence on three coacting factors (Fig. 6.a): the average basal phosphatase and kinase content *µ*_*N*_ and *µ*_*K*_, directly modulated by FK506 concentrations, and the stimulus induced quota *N*_*s*_, determined by FK506 and the stimulus features. The emerging average plasticity behaviour, in particular of the stimulated spine, is determined by specific values of these parameters, and, importantly, we see that there is a whole set of different values leading to no change in respect to the control condition (Fig. 6.k, gray surface). Moreover, due to the continuity of the change in potentiation with respect to these parameters, lower dosages of FK506 (1*µM*) would not substantially alter the basal/induced ratios, leading to statistically non-significant effects on synaptic plasticity [12].

In summary, our model is able to quantitatively characterize the interplay between the basal synaptic protein distributions and the molecular action of a synaptic stimulus, correctly reproducing the effect of catalyst inhibitors on synaptic plasticity. Moreover, it is able to provide a unified interpretation for the bidirectional effects on synaptic potentiation observed for the calcineurin inhibitor FK506, giving a clear interpretation of its dose-dependent effect.

## Discussion

We present a unified modeling framework describing the multi-spine plasticity occurring at the minute-to-hour timescale. We consider a molecular push and pull dynamic giving rise to variable plasticity outcomes and creating a distinct spatial footprint along a dendrite. We obtain interpretable driving equations from two fundamental biophysical processes, compatible with these spatial and temporal scales [7, 35, 36, 40, 41]: molecular diffusion and phosphorylation. Previous studies have followed a similar approach [9, 48, 49], with a notable example represented by [50], where the authors show how the general principle of resource-sharing can account for non-linear synaptic properties like multiplicative scaling and runaway dynamics prevention. Our model is able to rigorously extend this framework and endow it (1) with explicit biochemical meaning and (2) a clear spatial structure, both crucial to meaningfully investigate the principles driving synaptic plasticity.

This specification is crucial to understanding how an induction protocol impacts the plasticity dynamics of a multi-spine system. It allows, for example, for a clear differentiation between a “passive” depression induced by competition for resources [9, 50] and the “active” depression, mediated by the activation of the phosphatase family *N* in both stimulated and neighbouring spines. This latter form, in particular, is directly linked to the Mexican-hat shaped plasticity profile observed for single stimulations [3], and derives from the higher activation spread of phosphatases in comparison to kinases [37, 51]. A strong confirmation of this hypothesis could come, among other possibilities, by investigating the heterosynaptic plasticity induced by two close stimuli as a function of their distance (as in Fig. 5.d-g).

The specific features of the induction protocol are not, however, the only factors determining the plasticity response. In fact, our model strongly suggests that they determine only the average behaviour of a much more diversified and variable dynamic landscape [2, 7, 9, 12, 22]. We posit that a considerable portion of this variability is encoded in the synaptic basal catalytic distributions. A substantial corpus of modeling work has characterized the features and the origins of synaptic statistical properties, focusing in particular on what appears to be a generalized compatibility of a number of synaptic quantities with a log-normal distribution [4]. Several mechanisms have been proposed to explain this observation, ranging from the most fundamental multiplicative noise [4], to more sophisticated models [52] grounded on general stochastic processes [6, 53], or local binding mechanisms [5]. Despite providing an accurate characterization of the problem, these models often do not provide an immediate mapping between their driving parameters and the biochemical machinery underlying synaptic distributions. Moreover, an assumption on which all of this models implicitly rely is that dendritic spines represent a *statistical ensemble*, i.e., they can be considered a different instantiation of the same, stationary random process. By taking a different approach and constraining our model to depend only on elementary molecular dynamics, we are able to propose a different hypothesis: log-normal compatibility could originate spontaneously as the result of the elementary diffusive dynamics of synaptic proteins. This hypothesis is not only able to directly link the spine-size log-normality to the underlying catalytic log-normality, but is also able to avoid the ensemble assumption. Moreover, it is able to provide a robust, minimal mechanism for the observations in [11, 42], where log-normal compatible synaptic distributions are shown to emerge independently of neuronal activity and, therefore, are potentially not driven by an information encoding optimality principle.

The statistical characterization of the synaptic catalysts can be translated by the model to a probabilistic description of the synaptic response to plasticity induction. Several experimental and theoretical works have observed an inverse relationship between synaptic size (or weight) and its tendency to potentiate when stimulated, under a variety of plasticity protocols [2, 6, 12, 20, 52]. In some instances, this inverse relationship is not constrained to a specific plasticity direction, with the same induction being able to elicit both depression and potentiation, depending on the initial synaptic strength [54]. Our model is able to support this observation, in strong agreement with the power-law dependence proposed in [7]. Following this, we put forward the hypothesis that the synaptic response profile is quantitatively related to the calcium-induced catalytic dynamics occurring at every spine, and directly linked to the kinase-to-phosphatase ratio before and after stimulation. For small spines, starting from a low 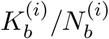, this ratio would on average increase, while the opposite would happen for large spines. Lastly, a class of intermediate spines, the size of which would depend on the stimulation features, would not show a change in its catalytic ratio, and therefore would appear to not respond to the induction protocol (as reported in, e.g. [54]). This mechanism is in line with the observations in [13], where partial inhibition of Protein Phosphatase 1 (one component of the model’s phosphatase family *N*) induces an overall shift towards potentiation, switching the response to a 10*Hz* simulus protocol from LTD to LTP. It is important to remark that our model, despite being able to predict the existence of potentiation in very small spines undergoing LTD-inducing stimulation [2], does show a lower quantitative predictive power in this latter case. This discrepancy could arise from several reasons, one of the most likely being the difference in biochemical pathways leading to potentiation or depression [14, 24]. Despite characterizing two possible general depression mechanisms (passive resource subtraction and active heterosynaptic phosphatase induction), the model is ultimately fitted on data obtained under a potentiation protocol. The emerging predictions could, therefore, not be optimal for describing dynamics mediated by other types of receptors like, for example, metabotropic glutamate receptors [14].

One final problem our model allows us to explore is the dependence of synaptic plasticity on catalyst perturbation and, in particular, on the differential block of calcineurin. The effect of FK506 on synaptic potentiation has been studied under a plethora of experimental conditions ([37, 55, 56] and references therein) with observations supporting both a facilitating and a hindering action. Multiple reasons have been proposed for the contradictory nature of these results, focusing strongly on the qualitative differences in the evoked calcium second-messenger cascade [57, 58] under different experimental conditions.

Our model suggests the possibility that these qualitatively different results could emerge from a quantitative feature, i.e, a degeneracy in the phospho-deposphatization dynamics with respect to their driving parameters. This degeneracy corresponds to the invariance of the observed synaptic potentiation when a change in the amounts of initial and stimulus-induced catalyst occurs. It has been shown that FK506 impacts both basal activity of CaMKII and calcineurin [37, 46], as well as the efficacy of the newly activated calcineurin quota in response to a plasticity protocol. These three components correspond directly to three model parameters (*µ*_*N*_, *µ*_*K*_ and *N*_*s*_) and, consequently, to three degrees of freedom. We show that this 3-dimensional space is foliated by the smooth maximal potentiation isosurfaces an average spine can show after stimulation. Different experimental conditions would correspond to different points in this space, belonging to different surfaces and, consequently, different degrees of synaptic potentiation. Moreover, the dose-dependent effect of FK506 would correspond to a smooth line traversing these isosurfaces. In order to characterize this line, however, a precise titration of the impact of FK506 on basal and newly activated CaMKII and CaN is necessary.

In this current study, we have largely treated spines and synapses as interchangeable. Both spines [59] and synapses have been shown to have a log-normal distribution as well as many other aspects of brain function (see [4] for a detailed review). While many aspects of spine volume correlate well with general synaptic structure and function, the spatial organization of a group of spines undergoing plasticity is not as well-studied. Typically, experimenters stimulate bundles of axons, which then synapse across the dendritic arbor of the post-synaptic neuron, onto multiple dendritic branches. In these cases (unless a single axon is being stimulated), multiple synapses sharing the same dendrite will likely be simultaneously active. Additionally, single axons have been demonstrated to make multiple clustered synaptic contacts onto individual dendrites in many cell types, including human cortex [60– 62], increasing the likelihood of simultaneous plasticity and competition for resources between spines sharing the same dendritic stretch. In all three spatial modalities, postsynaptic structures will have to compete for resources and be able to influence their neighbours.

In summary, we propose a model describing multi-spine plasticity and derive its equations from a minimal, although representative, set of biochemical interactions. We use this model to show that homo- and heterosynaptic plasticity can be jointly described in a multi-spine framework driven by two fundamental biochemical processes, i.e., phosphorylation and diffusion. By characterizing the basal synaptic catalytic distributions, we are able to put forward an alternative hypothesis for synaptic size log-normality, and to propose a mechanistic interpretation of the power-law-shaped dependence on initial size that spines show when undergoing potentiation. These insights can be used in future work to guide experimental design and interpretation, a well as provide vital theoretical constraints for the optimization of models including a more complex set of biochemical processes and spatio-temporal scales.

## Author information

J.P. and T.T. designed the study; T.E.C. carried out the glutamate uncaging expeirments; S.O.R. provided the pre- and post-synaptic protein quantification data; J.P. and M.F.E. analyzed the data; J.P. designed the model in collaboration with M.F.E., and optimized the parametrization with input from D.P and J.P.H.; J.P., M.F.E., T.E.C. and D.P. with input from all authors wrote the version of manuscript; All the authors contributed editing and writing.

## Acknowledgements

This research was supported by the University of Bonn Medical Centre, the University of Mainz Medical Centre, RIKEN Centre for Brain Science, OIST, JSPS Core-to-Core Programme (JPJSCCA20220007 to Y.G.), and by add-on fellowships of the Joachim Herz Stiftung (J.P., M.F.E.). We deeply thank Dr. Elba Raimúndez for the fruitful discussions and her valuable assistance with the model fitting. This project has received funding from the European Research Council (ERC) under the European Union’s Horizon 2020 research and innovation program (“MolDynForSyn”, grant agreement no. 945700 to T.T.) and was supported by the German Research Foundation via CRC 1286 (T.T., S.O.R.). This work was supported by the Open Access Publication Fund of the University of Bonn.

## Methods

### Experimental methods

#### Organotypic hippocampal slices

Slices were prepared as previously reported [63]. Briefly, hippocampi were isolated from P6-7 Wistar rat pups and cut into 350 *µ*m slices on a McIlwain tissue chopper. These were then cultured on cell culture inserts for 14-18 days in an incubator set to 35^◦^C in 5% CO2. The culture medium contained 50% Minimum Essential Medium (MEM, Thermo Fisher Scientific), 25% horse serum (Thermo Fisher Scientific), 23% Earle’s Balanced Salt Solution, and 36 mM D-glucose. For imaging experiments, slices were transferred to the microscope and perfused with room temperature aCSF containing (in mM) 125 NaCl, 2.5 KCl, 26 NaHCO3, 1.25 NaH2PO4, 20 glucose, 2 CaCl2, and 4 MNI-glutamate (Tocris). aCSF was continually bubbled with carbogen containing 95% oxygen and 5% carbon dioxide. In some experiments, calcineurin was inhibited with FK506 (2 *µ*M, Tocris). All animal experiments were approved by the RIKEN Animal Experiments Committee and performed in accordance with RIKEN rules and guidelines [Animal Experiment Plan Approval no. W2021-2-015(3)].

#### Transfection and imaging

Single neurons were biolistically transfected with a plasmid coding for EGFP using a Helios gene gun (Bio-Rad), and used for experiments 48–96 hours later. Imaging was performed on a Zeiss LSM 780 confocal laser scanning microscope, and all data were analysed offline. Experiments were limited to apical oblique dendrites of CA1 neurons.

#### Dendritic spine imaging and stimulation

Single dendrites were imaged for ≈ 1 hour, in which z stacks of the target dendrite were collected at every time point. Imaging was performed through a 63*×* objective, and each z step was 512 *×* 512 pixels, with 4*×*digital zoom, for a final frame size of 33.7 *µ*m. The z step was 0.5 *µ*m. MNI-glutamate was photolyzed at target spines using a custom-written macro controlling a 720 nm laser. Medium-sized mushroom spines were preferentially targeted for uncaging. The uncaging laser was positioned 0.5 *µ*m from the spine head, and used to deliver a train of laser pulses (60 pulses at 1 Hz, each pulse lasting 4 msec). For groups of stimulated spines, laser pulses were delivered in a quasi-simultaneous fashion in sequence, in which the first spine received a pulse of glutamate (4 msec), which was followed by a short delay (*<* 3 msec) as the system repositioned the laser to the next spine. This was repeated for all spines in the target cluster, and the sequence was repeated at 1 Hz for 60 cycles. For additional methods and details see [9].

### Experimental data analysis

This work relies on imaging datasets published in three previous works. We briefly describe the datasets and the respective analyses in the next paragraphs.

#### Model optimization and testing

We obtained the dataset published in [9], and integrated it with an additional experiment, where glutamate uncaging is performed at 5 closely located dendritic spines in control conditions, following the same experimental procedure (reported in the previous section of the Methods). Conducting image analysis in line with what was done in [9], we utilized the experiments conducted in control conditions to fit the optimal model parameters (1, 3, 5, 7 stimulation protocols). We utilized a different control experiment (protocol with 7 distributed stimulations) as a test to evaluate the predictive power of the model (Fig. 5.a). Finally, we utilized the data obtained with the application of FK506 (7 clustered stimulations) to investigate how phosphatase inhibition impacted the model’s behaviour (Fig. 6a-d). The synaptic fluorescence data deriving from these experiments was binned spatially and across different repetitions of the same protocol, following the procedure reported in detail in the Optimization section (Definition of the target values) in the Supplementary information.

#### Post-synaptic protein statistical analysis

We obtained the dataset presented in [8], where simultaneous quantification of synaptic size and protein content were performed via super-resolution microscopy in cultured hippocampal neurons. We analyzed the derived image dataset in line with the SpyDen pipeline [64] applied in [9], investigating the log-normality of the distributions across spines of plasticity-related proteins (Fig. 3e-g, Fig. S12).

#### Pre-synaptic protein statistical analysis

We obtained the dataset presented in [43], where presynaptic protein quantification was conducted in the cellular layers of the cortex and the cerebellum of adult rats. After synaptic bouton purification, electron microscopy, fluorescence microscopy, and quantitative immunoblotting were used to estimate the copy numbers of 64 different proteins per synaptosome. We analyzed the log-normal compatibility of the derived distributions (Fig. S13 by fitting a Gaussian curve to the logarithm of the protein copy numbers, and evaluating the fit relative root mean squared error (RRMSE). As a compatibility threshold, we considered a value of RRMSE *<* 20%.

### Synaptic size equation

Here, we give an overview of the steps we carried out to obtain our equation for synaptic sizes. The full derivation with the necessary mathematical details is discussed thoroughly in the Supplementary Information.

The main model differential system describes the *concentration* dynamics of the considered protein resources throughout the dendrite and the spines, together with the related changes in synaptic sizes. We are, therefore, interested in finding a simple equation that explicitly tells the size value of each of the spines located on the dendrite for a given value of time *t*. To achieve this equation, we exploit two main observations.

First, we expect that for the considered time span (roughly 1 hour), the net influx of new (unphosphorylated) proteins into the dendritic stretch can be neglected, due to its overall low value [65, 66]. This fact translates to no-flux boundary conditions for our system, which in turn lead to the conservation in time of the total amount of available synaptic resources ℛ

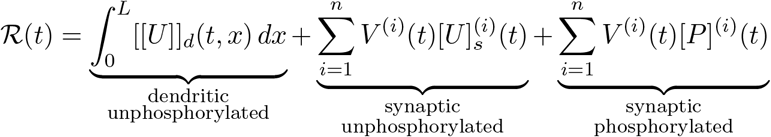

Second, we notice that multiple timescales are involved in the process. In particular, the (de)phosphorylation of *U*_*s*_ and *P*, as well as the diffusion of *U*_*d*_, can be considered fast compared to the variation timescales of the activated catalysts *K* and *N* [36, 67, 68]. Conversely, the timescale at which the amount of activated catalysts decay, corresponds to the timescale at which the experiment used in this work was conducted (tens of minutes, [9, 36]), and can be considered the leading timescale of the considered size dynamics. This allows us to resort to a quasi-steady-state approximation of the full differential model, which, together with the mass constraint for ℛ leads to the general equation for synaptic sizes

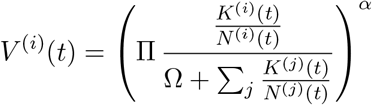

where we have assumed a power-law dependence *V* ^(*i*)^ ∝ (*P* ^(*i*)^)^*α*^of the synaptic sizes on the synaptic phosphorylated resource. This assumption derives from the structural role of *P* ^(*i*)^ in building the synaptic scaffolding, which can be well approximated with a power-law function. As one of the main components of *P* ^(*i*)^ is represented by actin, which directly correlates with synaptic size [49, 69], we choose to consider *α* = 1 (reducing as much as possible the free parametrization in accordance with the available experimental precision), obtaining the final equation

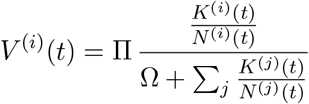

Importantly, fixing the value of *α* does not structurally impact the validity of these derivations, as no singularities are present for *α* = 1.

### Simulating a plasticity experiment

Once the set of optimal parameters has been found (the optimization procedure is described in detail in the relevant Supplementary Information section), we can use it to simulate a plasticity induction experiment following three steps:

1. we instantiate a dendrite with a given number of spines, and for each of the spines we draw a value for 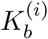 and 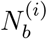 from their inferred bivariate log-normal distribution (extensively described in the Results section);
2. we select which spines will be targeted for plasticity induction. This selection can be either done arbitrarily or, for model testing, in accordance with the stimulations proposed in an experimental work;
3. using our model (2), we compute for each spine position *x*_*i*_ and time *t* the value of the synaptic size *V* ^(*i*)^(*t*).

For a given selection of stimulated spines, we repeat these steps multiple times, and express the final results in terms of the summary statistics emerging from the bivariate log-normal distribution of 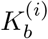and *N*_*b*_(*i*).

Importantly, since the model has been trained on the data deriving from [9], our best parametrization describes plasticity occurring in response to that specific stimulation protocol. Generalizing this model to different stimulation protocols or different neuronal populations can be achieved by changing the stimulus-related parametrization (*K*_*s*_, *N*_*s*_, *τ*_*K*_, *τ*_*N*_, *σ*_*K*_, and *σ*_*N*_), or the basal catalytic one (which is intrinsic to the neurons in consideration).

### Relation between catalyst and spine size log-normality

In the derivation of the synaptic size equation (reported in detail in the Supplementary Information) quasi-steady state approximation leads to the intermediate relation

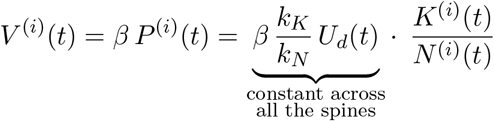

as 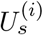 has balanced out with the dendritic quota *U*_d_, which itself is constant along the dendrite. For a fixed time *t*, the probability density of *P* ^(*i*)^ will therefore be equal to the ratio of the probability densities of *K*^(*i*)^ and *N* ^(*i*)^. Having found that these two densities are jointly log-normal, their ratio, and consequently synaptic sizes, follow a log-normal distribution.

### Non-responder synapse equation

We start by considering the synaptic size equation (2), and rewrite it for convenience as

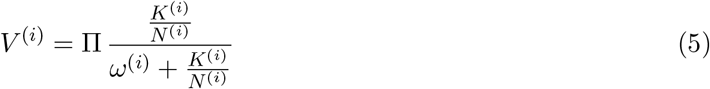

where 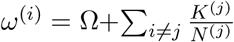 includes the heterosynaptic competitive portion of the system. As described in the previous sections, each spine starts from some basal catalytic values 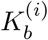 and 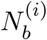, and a stimulus acts by modifying these values by some amount Δ_*K*_ and Δ_*N*_ depending on various factors (e.g., distance from the considered spine, time elapsed from induction). Importantly, this action takes place in every spine in the system, so that *ω*^(*i*)^ is also modified by some amount Δ_*ω*_. We focus our attention on stimulated spines, i.e. spines where Δ_*K*_ = *K*_*s*_ and Δ_*N*_ = *N*_*s*_. Importantly, calling 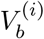 and 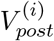 the basal and the post-induction sizes of a spine in consideration, we are interested in finding the relation between 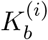 and 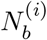 satisfying 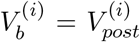. By applying (5) to both sides of the equation, one obtains

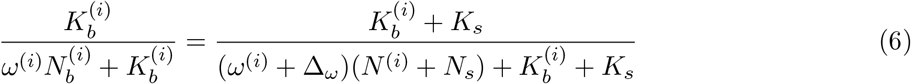

which can then be solved to obtain the final solution

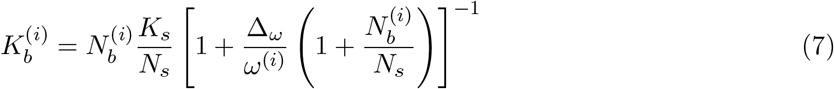

describing the curve in the *K* and *N* phase plane corresponding to the spines that will not change size in response to a stimulus inducing *K*_*s*_ and *N*_*s*_ amounts. This expression is stochastic, as for a fixed value of 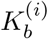 and 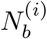, the heterosynaptic factor *ω*^(*i*)^, as well as it variations, depend on the specific dendritic instance. We explore this stochasticity by running multiple simulations using our optimal parameter set, and find that the value of Δ_*ω*_*/ω*^(*i*)^ is extremely small in all cases. This allows to approximate (7) to the linear equation 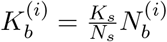, showing that the non-responding spines are exactly those with a basal catalytic ratio 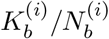 equal to the stimulus catalytic ratio *K*_*s*_*/N*_*s*_..

