## Supplementary information for "Push-and-pull protein dynamics leads to log-normal synaptic sizes and probabilistic multi-spine plasticity"

### Derivation of the closed-form synaptic size equation

**Fundamental definitions** For our derivation, we follow the established ODE/PDE framework [1]. We start with the following definitions:

- the *dendrite* as the set $D = [0, L] \subset \mathbb{R}$ with length $L$, with $n$ spines located at positions $x_i \in D, i = 1, ..., n$;
- the *time interval* throughout which our process takes place is the real valued domain $T \subset \mathbb{R}$;
- the *dendritic unphosphorylated resource* is as positive function $U_d : D \times T \to \mathbb{R}^+$;
- the *synaptic unphosphorylated resource* is defined as $n$ positive functions, $U_s^{(i)} : T \to \mathbb{R}^+$, each describing the amount of the resource in the spine located at position $x_i$ for each time $t$. Notice that $U_d$ and $U_s$ represent the same molecular species (unphosphorylated resource) but describe its behaviour in different spatial compartments (dendrite and spines);
- the remaining synaptic quantities, i.e. *activated kinases, activated phosphatases, phosphorylated synaptic resource, and synaptic size* are defined in line with $U_s$, and identify the four function families $K^{(i)}, N^{(i)}$, $P^{(i)}$, and $V^{(i)}$;
- the synaptic (volumetric) concentration of a quantity $X^{(i)}$ is defined as $[X]^{(i)} = X^{(i)}/V^{(i)}$
- $\ell$ is a unit of dendritic length, necessary for the correct computation of dendritic concentrations;
- the dendritic (linear) concentration of a quantity $Y$ is defined as $[[Y]]^{(i)} = Y^{(i)}/\ell$

We assume the functions defined above present a sufficient degree of smoothness to undergo the usual differentiation, as they represent classical, well-behaved physical quantities. In virtue of these definitions, the driving equations of the diffusion-phosphorylation process unwinding throughout a linear dendritic branch and its dynamically evolving spines can be written as

$$\begin{cases} \dfrac{d[P]^{(i)}}{dt} = k_K[K]^{(i)}\,[U]_s^{(i)} - k_N[N]^{(i)}\,[P]^{(i)} - \dfrac{[P]^{(i)}}{V^{(i)}}\dfrac{dV^{(i)}}{dt} \\ \dfrac{d[U]_s^{(i)}}{dt} = \dfrac{k_{in}\ell}{V^{(i)}}\,[[U]]_d - k_{out}[U]_s^{(i)} - k_K[K]^{(i)}\,[U]_s^{(i)} + k_N[N]^{(i)}\,[P]^{(i)} - \dfrac{[U]_s^{(i)}}{V^{(i)}}\dfrac{dV^{(i)}}{dt} \\ \dfrac{\partial[[U]]_d}{\partial t} = D_U\,\dfrac{\partial^2[[U]]_d}{\partial x^2} - \sum_{i=1}^{n}\delta(x - x_i)\left[k_{in}\ell\,[[U]]_d - k_{out}V^{(i)}\,[U]_s^{(i)}\right] \end{cases} \tag{S.1}$$

where

- $k_K$ and $k_N$ are the phosphorylation rates of kinases and phosphatases, respectively;
- $k_{in}$ and $k_{out}$ are the exchange rates between spine and dendrite;
- $D_U$ is the diffusion coefficient for the unphosphorylated resource;

- $\delta(x)$ is the Dirac-delta function.

The size variation of the involved spines translates to the appearance of the volume derivative terms directly from the derivative of the ratio. For some general species $X$, we have, indeed, that

$$\frac{d}{dt}[X]^{(i)} = \frac{d}{dt}\left(\frac{X^{(i)}}{V^{(i)}}\right) = \frac{1}{V^{(i)}}\frac{dX^{(i)}}{dt} - \frac{[X]^{(i)}}{V^{(i)}}\frac{dV^{(i)}}{dt} \tag{S.2}$$

with $dX^{(i)}/dt$ corresponding to the net flux of incoming our outgoing molecules $X^{(i)}$. Importantly, the positivity of each of the involved quantities (evolving species and driving constants) ensure that the dynamics take place only in the positive region of the state space (negative terms vanish when the compartment they are subtracting from reaches 0 value).

Since the unbiased influx and outflux of protein resources into the dendrite can be considered extremely low during the time span of the studied phenomenon ([2], $T \sim 1$hour) we associate to the system (S.1) no-flux boundary conditions at the dendrite extremities:

$$\frac{\partial[[U]]_d}{\partial x}(t,0) = \frac{\partial[[U]]_d}{\partial x}(t,L) = 0 \tag{S.3}$$

**Working assumptions** before proceeding with the derivation, we introduce some working assumptions:

1. the dependence between synaptic volume and the amount of underlying structural proteins is active topic of experimental and computational research, and there is currently no universally accepted equation that describes the relationship between $V^{(i)}$ and $P^{(i)}$. It can be agreed on, however, that the phosphorylated resource is indeed used as a structural block of the synaptic structure, acting either as the scaffolding (radial or superficial) of a rigid spine, or as a filling, "inflating" an elastic spine. All these implementations can be, in first approximation, modeled as a power-law dependence of $V^{(i)}$ on $P^{(i)}$

$$V^{(i)} := \beta P^{(i)\,\alpha}$$

   with $\beta$ being a dimensional constant;

2. the dendritic shape and volumes do not change significantly at the considered timescale;

3. $k_{in}$ and $k_{out}$ are the exchange rates across the spine neck. To the best of our knowledge, there is no consensus on the existence of a bias in this exchange (which is reasonable, given the symmetry of the neck). Therefore, in our derivation, we will assume that $k_{in} = k_{out} := k_{ex}$;

4. in accordance with experimental observations, none of the involved quantities are ever equal to 0 - they can be small but never null. Loss of dendritic spines has been observed (e.g. pruning), but it is mediated by a slower molecular processes, potentially involving mRNA translation and not only fast reversible plasticity changes. Therefore, we do not consider spine removal in our model;

5. **quasi-steady-state (QSS) assumption:** the phosphorylation and diffusion processes are fast with respect to the changes of $K^{(i)}$ and $N^{(i)}$ in time. This allows us to consider the catalyst concentration as constant and impose a steady-state condition for the remainder of the system. Ultimately, we will recover the dependence on time by reintroducing the evolution of $K^{(i)}$ and $N^{(i)}$ in the steady-state solution.

**Conservation of total resource amount** We are now interested in finding a closed-form expression for the evolution of the synaptic sized $V^{(i)}$ in time.

First, we start by noticing that the no-flux boundary conditions lead to the conservation of the total amount of resources throughout time. By defining this resource amount as $\mathcal{R}(t)$

$$\mathcal{R}(t) := \underbrace{\int_0^L [[U]]_d(t,x)\,dx}_{\substack{\text{dendritic}\\ \text{unphosphorylated}}} + \underbrace{\sum_{i=1}^n V^{(i)}(t)[U]_s^{(i)}(t)}_{\substack{\text{synaptic}\\ \text{unphosphorylated}}} + \underbrace{\sum_{i=1}^n V^{(i)}(t)[P]^{(i)}(t)}_{\substack{\text{synaptic}\\ \text{phosphorylated}}} \tag{S.4}$$

we can verify that its derivative in time is null

$$\begin{aligned}
\frac{d\mathcal{R}}{dt} =& \frac{d}{dt}\left\{\int_0^L [[U]]_d\,dx + \sum_i V^{(i)}\left[[U]_s^{(i)} + [P]^{(i)}\right]\right\} = \\
=& \int_0^L \frac{\partial [[U]]_d}{\partial t}\,dx + \sum_i \frac{dV^{(i)}}{dt}\left[[U]_s^{(i)} + [P]^{(i)}\right] + \sum_i V^{(i)}\left[\frac{d[U]_s^{(i)}}{dt} + \frac{d[P]^{(i)}}{dt}\right] = \\
=& \left.\frac{\partial [[U]]_d}{\partial x}\right|_0^L = 0
\end{aligned}$$

where the last step is given by the evaluation of $[[U]]]_d$ at the boundary conditions locations. Therefore, $\mathcal{R}(t) = \mathcal{R}$ is constant in time.

**QSS approximation** As previously introduced, the concentration variations depend both on the influx and outflux of molecules and the variation $V^{(i)}$. By inserting (S.2) into the left-hand-side of our model (S.1), we can obtain the equations driving the change in time of the absolute quantities of $U$ and $P$

$$\begin{cases}
\dfrac{dP^{(i)}}{dt} = \dfrac{k_K K^{(i)}\,U_s^{(i)} - k_N N^{(i)}\,P^{(i)}}{V^{(i)}} \\
\dfrac{dU_s^{(i)}}{dt} = k_{in} U_d - k_{out} U_s^{(i)} - \dfrac{k_K K^{(i)}\,U_s^{(i)} - k_N N^{(i)}\,P^{(i)}}{V^{(i)}} \\
\dfrac{\partial U_d}{\partial t} = D_U\,\dfrac{\partial^2 U_d}{\partial x^2} - \ell \displaystyle\sum_{i=1}^n \delta(x - x_i)\left[k_{in}\,U_d - k_{out} U_s^{(i)}\right]
\end{cases} \tag{S.5}$$

We are now interested in the solution for the steady state

$$(\dot{P}^{(i)}, \dot{U}_s^{(i)}, \partial_t U_d) = (0, 0, 0)$$

We mention that the only singularity that could compromise the dynamics of this system appears for $V^(i) = 0$, which has been excluded in the working assumptions. By proceeding and imposing the steady state conditions, we get

$$\begin{cases} P^{(i)} = \dfrac{k_K\, K^{(i)}}{k_N\, N^{(i)}} U_s^{(i)} \\ U_s^{(i)} = U_d(x_i) \\ \partial_x^2 U_d = 0 \end{cases} \tag{S.6}$$

The third equation (one-dimensional Laplace equation) coupled with no-flux boundary conditions has constant solution in space $U_d(x) = \bar{U}_d$. This result, together with the other relationships of (S.6) can be immediately replaced in the definition of $\mathcal{R}$ (S.4), yielding the relationship

$$\mathcal{R} = \frac{\bar{U}_d}{\ell} L + \bar{U}_d \sum_{i=1}^{n} \left[ 1 + \frac{k_K\, K^{(i)}}{k_N\, N^{(i)}} \right]$$

which leads to the closed expression for $\bar{U}_d$ only in terms of constant parameters

$$\bar{U}_d = \frac{\mathcal{R}}{\frac{L}{\ell} + n + \sum_i \frac{k_K K^{(i)}}{k_N N^{(i)}}}$$

Notice that $L/\ell$ is just the pure number describing the length of the dendrite expressed in the unit of measure $\ell$. What is left to do is to substitute this expression into the QSS expression for $P^{(i)}$ from (S.6), obtaining

$$P^{(i)} = \Pi \frac{\frac{k_K\, K^{(i)}}{k_N\, N^{(i)}}}{\frac{L}{\ell} + n + \sum_j \frac{k_K\, K^{(j)}}{k_N\, N^{(j)}}}$$

After defining

1. the linear synaptic density as $\lambda = n\ell/L$
2. the parametric constant as $\Omega = \frac{k_N L(1+\lambda)}{k_K \ell}$;
3. the conserved, maximally achievable size $\Pi = \beta\, \mathcal{R}$

and recalling the relation $V^{(i)} = \beta P^{(i)\,\alpha}$ we obtain the closed-form expression for the synaptic sizes

$$V^{(i)}(t) = \left( \Pi \frac{\frac{K^{(i)}(t)}{N^{(i)}(t)}}{\Omega + \sum_j \frac{K^{(j)}(t)}{N^{(j)}(t)}} \right)^{\alpha}$$

where we have made explicit again the slow dependence on time of the catalytic ratio $K^{(i)}/N^{(i)}$. Ultimately assuming, in accordance with the observed literature, direct proportionality between $P^{(i)}$ and $V^{(i)}$ ($\alpha = 1$) we obtain the final expression for the synaptic sizes.

### Optimization

**Structural identifiability** Our model describes the evolution of each spine on a dendrite subject to a specific plasticity induction protocol. Consider now a scenario where $m = 1, ..., M$ different induction protocols have been carried out on different neurons. Each of these scenarios entails different stimulation locations and different dendrites, with different geometric features, different quantities of available resources, and different basal catalytic conditions.

Indicating with $\chi^{(i)}$ the catalytic ratio at the $i$-th spine after a stimulus has been applied at time 0

$$\chi_m^{(i)}(t) = \frac{K_{b,m}^{(i)} + \Theta(t)\, K_s\, e^{-\frac{t}{\tau_K}} \sum_{\bar{x}\in\mathcal{X}_m} e^{-\frac{(x_i-\bar{x})^2}{\sigma_K^2}}}{N_{b,m}^{(i)} + \Theta(t)\, N_s\, e^{-\frac{t}{\tau_N}} \sum_{\bar{x}\in\mathcal{X}_m} e^{-\frac{(x_i-\bar{x})^2}{\sigma_N^2}}} \tag{S.7}$$

for each condition $m$ our model is formulated as

$$\begin{cases} V_m^{(i)}(t) = \Pi_m \dfrac{\chi_m^{(i)}(t)}{\Omega_m + \sum_{j=1}^{n_m} \chi_m^{(j)}(t)} \qquad m = 1, ..., M \end{cases} \tag{S.8}$$

where, assuming that the stimulation protocol happens at $t = 0$

The number $n_m$ of observed spines for each protocol, and the locations of the stimulations ($\mathcal{X}_m = \{\bar{x}_{1,m}, ..., \bar{x}_{1,m}\}$) are known a priori from the experimental procedure. The other parameters used in the equations, however, are not known and need to be recovered via data fitting. Conceptually, we can distinguish these parameters into two groups:

1. *global parameters*, driving the plasticity dynamics independently of the specific experimental realization, and depending on the stimulation features. These are $K_s$, $N_s$m, $\tau_K, \tau_N, \sigma_K$, and $\sigma_N$;
2. *specific parameters* which depend on a specific experimental setup. These are the spine initial conditions ($K_{b,m}^{(i)}, N_{b,m}^{(i)}$), the available resources $\Pi_m$, and the dendritic parametric constant $\Omega_m$. This parameter category is indexed by the protocol index $m$.

| **Global parameters** | | **Number** |
|---|---|---|
| Timescales | $\tau_N$, $\tau_K$ | 2 |
| Length scales | $\sigma_K$, $\sigma_N$ | 2 |
| Stimulus contribution | $K_s$, $N_s$ | 2 |

| **Specific parameters** | | **Number** |
|---|---|---|
| Total resources | $\Pi_m$ | $M$ |
| Dendritic geometric factor | $\Omega_m$ | $M$ |
| Initial kinases per spine | $K_{b,m}^{(i)}$ | $\sum_{m=1}^{M} n_m$ |
| Initial phosphatase per spine | $N_{b,m}^{(i)}$ | $\sum_{m=1}^{M} n_m$ |

Table S1: Model parameters necessary for the simultaneous optimization of $M$ protocols with $n_m$ spines each. The specific parameters are indexed by the protocol index $m$, and describe quantities that can differ between experiments, like the total available resources in the dendrite, the geometry of the dendrite and the synaptic catalyst distributions at stimulus time.

The total number of parameters that have to be inferred when fitting data arising from $M$ experimental protocols amounts to $6 + 2\,M + 2\,\sum_m n_m$ (ref. Tab. S1). It is not possible, however, to fit all these parameters together due to the presence of structural non-identifiability in this native form of the model. This fact arises from the rescaling invariance for $V_m^{(i)}$ and $\chi_m^{(i)}$ in (S.8) and (S.7):

$$V_m^{(i)}\left(\Pi_m, \chi_m^{(1)}, ..., \chi_m^{(n_m)}, \Omega_m\right) = V_m^{(i)}\left(\Pi_m, s\,\chi_m^{(1)}, ..., s\,\chi_m^{(n_m)}, s\,\Omega_m\right), \quad s \in \mathbb{R}$$

$$\chi^{(i)}\left(t; K_{b,m}^{(i)}, K_s, N_{b,m}^{(i)}, N_s\right) = \chi^{(i)}\left(t; s\,K_{b,m}^{(i)}, s\,K_s, s\,N_{b,m}^{(i)}, s\,N_s\right), \quad s \in \mathbb{R}$$

This can be solved by introducing two additional constraints, one for each of the two degrees of freedom. To this end, we temporarily set the values of $\Omega_M$ and $N_{b,M}^{(n_M)}$ to 1 (one dendritic constant, and the basal amount of phosphatases of one spine), and express the remainder of the parameters with respect to them.

Defining, for practicality, the "fractionary index" symbols

$$\Omega_{m/M} = \frac{\Omega_m}{\Omega_M}, \quad K_{b,m/M}^{(i)} = \frac{K_{b,m}^{(i)}}{\Omega_M N_{b,M}^{(n_M)}}, \quad N_{b,m/M}^{(i)} = \frac{N_{b,m}^{(i)}}{N_{b,M}^{(n_M)}}, \quad K_{s/M} = \frac{K_s}{\Omega_M\, N_{b,M}^{(n_M)}}, \quad N_{s/M} = \frac{N_s}{N_{b,M}^{(n_M)}}$$

we can rewrite the problem equations (S.8)

$$
\begin{cases}
V_m^{(i)}(t) = \Pi_m \, \dfrac{\chi_m^{(i)}(t)}{\Omega_{m/M} + \sum_{j=1}^{n_m} \chi_m^{(j)}(t)} & m = 1, ..., M-1 \\
V_M^{(i)}(t) = \Pi_M \, \dfrac{\chi_M^{(i)}(t)}{1 + \sum_{j=1}^{n_M} \chi_M^{(j)}(t)} & m = M
\end{cases}
\tag{S.9}
$$

where

$$
\chi_m^{(i)}(t) = \frac{K_{b,m/M}^{(i)} + K_{s/M}\, e^{-\frac{t}{\tau_K}} \sum_{\bar{x}\in\mathcal{X}_m} e^{-\frac{(x_i-\bar{x})^2}{\sigma_K^2}}}{N_{b,m/M}^{(i)} + N_{s/M}\, e^{-\frac{t}{\tau_N}} \sum_{\bar{x}\in\mathcal{X}_m} e^{-\frac{(x_i-\bar{x})^2}{\sigma_N^2}}}, \qquad m = 1, ..., M-1
$$

$$
\chi_M^{(i)}(t) = \frac{K_{b,m/M}^{(i)} + K_{s/M}\, e^{-\frac{t}{\tau_K}} \sum_{\bar{x}\in\mathcal{X}_m} e^{-\frac{(x_i-\bar{x})^2}{\sigma_K^2}}}{1 + N_{s/M}\, e^{-\frac{t}{\tau_N}} \sum_{\bar{x}\in\mathcal{X}_m} e^{-\frac{(x_i-\bar{x})^2}{\sigma_N^2}}}, \qquad m = M
$$

With this reduced parametrization, the problem (S.9) is well posed. We can then proceed to define a biologically plausible value range for the free parameters (Tab. S2) and carry out the inference. As a final step, in accordance with the previous literature [3], we select a reasonable value for the two constraints $\Omega_M$ and $N_{b,M}^{(n_M)}$, and recover with respect to them the optimal values for the full optimization problem (S.8).

**Definition of the target values** The equations (S.8) describe, for every protocol $m = 1, ..., M$, the dynamics of the $n_m$ spines located on one dendrite. Since in our experimental data, each protocol is reproduced multiple times on different cells, we have to build a dataset containing one statistically representative dendrite for each protocol. We proceed as follows:

1. we select an experimental protocol $m$, uniquely defined by the stimulations $\mathcal{X}_m$ of stimulations provided;
2. **distance assignment:** for each cell, we assign to each synapse its distance value from the nearest stimulation. Importantly, if the spine is located between two stimulations, we assign it a negative distance value. We assign a distance value of 0 to the stimulated spines;
3. **average spine density:** for the model (S.8) to reproduce the studied dynamics, we have to consider the correct number of spines located on the dendrite. We infer this parameter by estimating the mean inter-spine distance across all the cells used for each protocol. Details and benchmarking of the estimation procedure are reported in "Inter-spine distance estimation" section of the Supplementary Information;
4. **binning:** for each protocol $m$ we generate a statistically representative dendrite with spines located regularly at the inferred inter-spine distance, and size values equal to the luminosity

| Parameter | Unit | BVs | Notes |
|---|---|---|---|
| $K_{b,m/M}^{(i)}$ | - | $0.002 - 0.12$ | Supposing 4% of CaM is active basally, and equally split between $K$ and $N$ [3–5] |
| $K_{s/M}$ | - | $0.07 - 3.71$ | Assuming that CaM is the limiting activation factor [3–6] |
| $N_{b,m/M}^{(i)}$ | - | $0.28 - 3.5$ | Using the same argument as in $K_{b,m/M}^{(i)}$ [3–5] |
| $N_{s/M}$ | - | $0.07 - 3.71$ | Assuming that calcineurin is the limiting factor [3–5] |
| $\tau_K$ | min | $0 - 100$ | Covering bulk, local and reciprocal kinase-effector activation timescales [5] |
| $\tau_N$ | min | $0 - 100$ | Range as in $\tau_K$ |
| $\sigma_K$ | $\mu m$ | $0 - 100$ | Wide parameter range |
| $\sigma_N$ | $\mu m$ | $0 - 100$ | Wide parameter range |
| $\Omega_{m/M}$ | - | $0.3 - 3.33$ | Observations in [3] and general dendritic statistics |

Table S2: Parameter boundary values (BVs) used in the optimization

average across different cells of the corresponding spatial bin. To avoid overlapping, we choose a bin with equal to the inferred inter-spine distance (Fig. S1).

In this fashion, we obtain a dataset consisting of 5 raw integrated density fields, one for each protocol, effectively describing the synaptic size evolution before and after plasticity induction. We use 4 of these datasets for model optimization ($1, 3, 5$, and 7 clustered stimulation protocols), and leave out one (7 distributed stimulation protocol) for model validation.

One final factor that needs to be considered before proceeding with the optimization is that the datasets obtained with this procedure contain a reduced amount of spines (Fig. S1.a,b). This would lead, during the fitting, to an underestimation of the factor $\sum_j \alpha^{(j)}(t)$, in the denominator of (S.8). To correctly account for all the spines in the experimental dataset, we introduce a differential weighting of the spines depending on their type (stimulated, inside of the stimulation cluster, and outside of the stimulation cluster), and compute the sum as

$$\sum_{j=1}^{n_m} \alpha^{(j)} \simeq |\mathcal{X}_m|\, \alpha^{(0)} + 2 \sum_{j \in \text{inside}} \alpha^{(j)} + (2 - |\mathcal{X}_m|) \sum_{j \in \text{outside}} \alpha^{(j)} \tag{S.10}$$

where $|\mathcal{X}_m|$ is the cardinality of $\mathcal{X}_m$, i.e. the number of stimulations for the protocol $m$.

**Parameter estimation** To perform parameter estimation, we used the maximum likelihood estimate (MLE) approach, finding the values of the model parameters $\theta$ that maximize the likelihood function $p(\theta|\mathcal{D})$ of the dataset $\mathcal{D}$ under some noise assumptions.

$$\theta_{MLE} = \arg\min_{\theta} \left[-\log p(\mathcal{D}|\theta)\right] \tag{S.11}$$

The dataset $\mathcal{D}$ consists of the binned synaptic size observations $y_m^{(i)}(t_j)$, where $(i)$ is the spatial index of the synaptic bin, $t_j$ is the discrete temporal coordinate of the observation, and $m$ is the protocol index, indicating from which of the protocols in [7] the observation comes from (for the optimization, we utilized the clustered 1, 3, 5, and 7 stimulations under control conditions).

Since our target derives from spatial binning, it has an associated Student t-distributed uncertainty, with degrees of freedom $d_m^{(i)}$ equal to the number of spines located inside the bin. We used this uncertainty as our noise model, preferring it to standard Gaussian noise for its better performance when optimizing models on outlier-rich data [8]. The resulting negative log-likelihood can be, therefore, written as

$$\mathcal{J}(\theta) = -\log p(\mathcal{D}|\theta) = \sum_{i,j,m} \frac{d_m^{(i)}+1}{2} \log \left\{ 1 + \frac{\left[y_m^{(i)}(t_j) - \hat{y}_m^{(i)}(t_j)\right]^2}{d_m^{(i)}} \right\} + C$$

where $\hat{y}_m^{(i)}(t_j)$ denotes the model prediction, and $C$ is a constant parameter deriving from the Student t-distribution renormalization term. In order to minimize $\mathcal{J}(\theta)$ (find the maximum likelihood estimate), we performed gradient-based optimization on the log-transformed parameters, using a multi-start strategy from 1200 different initial values. We implemented the model and $\mathcal{J}$ in PyTorch [9], and resorted to the Python Parameter Estimation Toolbox (pyPESTO) with the Fides optimizer to perform gradient minimization [10, 11].

The optimal fit-to-data is shown in Fig. S3, and a more thorough analysis of the optimization outcome is reported in the next section

**Reliability and uncertainty quantification** We assessed the quality of the optimization results by comparing the final objective function values for the different optimization runs (Fig. S2). As shown in the waterfall plot (Fig. S2.a), there are two wide local minima (LM1 and LM2) in the parameter space, with 437 and 656 converging runs, respectively. The minima show almost identical final criterion values ($fval_1 = 108.88$, $fval_2 = 108.96$), as well as very similar inferred values for the parameters. Interestingly, the only substantial difference seems to reside in the estimates for the decay constants $\tau_K$ and $\tau_N$, with LM2 converging on significantly higher values (Fig. S2.b).

Following standard practice, we considered the final parameters of the optimization run which reached the lowest objective function value as maximum likelihood estimates. The resulting fits to data, shown in Fig. S3, display remarkable qualitative accuracy, correctly reproducing both homosynaptic potentiation at $x^{(i)} = 0.00\,\mu m$ and heterosynaptic depression, when present. Quantitatively, the model achieves a 9.72% relative root mean squared error (RRMSE), with narrow and symmetric residual and relative residual distributions (Fig. S4). Remarkably, the same metrics are obtained using values for

$\tau_K$ and $\tau_N$ deriving from the best run of LM2. Given their better compatibility with the investigated timescales, we used these values in our simulations.

As a final step in the optimization evaluation, we estimated the posterior distributions of the inferred parameters. Given the very high dimensionality of the model's parameterization, we split this procedure into two parts, using two different approaches (a full Markov chain Monte Carlo sampling with a sufficiently high number of samples $> 100000$ resulted computationally prohibitive with the available equipment).

For the synaptic basal catalytic, we resorted to an ensemble-based uncertainty quantification focusing on the 437 optimization runs belonging to LM1. This revealed that the rescaled basal values $K^{(i)}_{b,m/M}$ and $N^{(i)}_{b,m/M}$ follow, with very few exceptions, a smooth, symmetric distribution, well contained within the chosen parameter bounds (Fig. S6). Visual analysis already hints at log-normal compatibility of these estimates.

Ultimately, we fixed the basal catalytic values to their MLE estimates, and performed a Markov chain Monte Carlo uncertainty quantification for the six remaining global parameters (the maximal catalytic activations $K_{s/M}$ and $N_{s/M}$, the timescales $\tau_K$ and $\tau_N$, and the spatial scales $\sigma_K$ and $\sigma_N$) using an adaptive parallel tempering algorithm [12]) with 10 chains and $10^5$ samples (Fig. S5. Interestingly, some degree of degeneracy seems to be present for the values of $\tau_K$ and $\tau_N$, compatibly with the two different convergence values shown by LM1 and LM2.

### Inter-spine distance estimation

We are interested in estimating the mean inter-synaptic distance ($MISD$) from the data describing a linear dendritic stretch. As stated in the Methods section, each spine is given a distance value corresponding to the distance from the closest stimulation. Moreover, this value is negative in case the considered spine lies inbetween different stimulations and positive otherwise.

We start by assuming that the linear spine density is homogeneous, i.e. it does not change with the absolute position along the dendrite at the considered length scales ($10 - 100\,\mu m$).

From the array of distances, we then consider only the positive values, deriving from the spines located outside the stimulation cluster; we then sort this array in ascending order. This sorted array now contains roughly double the amount of spines located on an average dendritic stretch, as what we have done is mapped two different stretches onto one. In order to estimate the average inter-spine distance, led by this heuristic, we subsample this array, taking every second distance value.

As a final step, we compute the differences between consecutive distances and use their average value as an estimate of the true inter-spine distance.

We quantify the goodness of this estimator using a Monte-Carlo sampling. We generate $N = 10^5$ different dendrites with 100 spines each. In accordance with [13], we use a Weibull distribution

$$f(x; c, s) = \frac{c}{s}\left(\frac{x}{s}\right)^{c-1} e^{-(x/s)^c} \tag{S.12}$$

with fixed scale and shape parameters ($s$ and $c$) to generate random inter-synaptic distances for each dendrite. We then carry out our estimation using the procedure described above, and evaluate its performance in terms of the mean relative error

$$MRE = \left\langle \frac{\overline{MISD}}{MISD} - 1 \right\rangle \tag{S.13}$$

where $\overline{MISD}$ is the estimated mean inter-spine distance and $MISD$ is the true theoretical value $MISD = s\Gamma(1 + 1/c)$. We quantify this metric on a set of different values of $c$ and $s$, focusing on parameter ranges giving rise to mean inter-spine distances between 1 and a few microns.

The results, reported in Fig. S7, show that the described estimator has a well-behaved, bell-shaped distribution, with an average bias of $\sim$ 3%, confined under a 30% error. We consider this error acceptable since it its absolute value ($\sim 0.8\,\mu m$) is comparable with the spatial resolution of the experimental setup used to collect the data.

### Stochastic simulation of a diffusive molecule in a dendritic stretch

Consider a one-dimensional dendrite of length $L$ with spines located at positions $x = x_i$, $i = 1, ..., N$. Consider now an abstract molecular family able to diffuse throughout the dendrite, entering and exiting the dendritic spines (with fixed size, for simplicity). Let now $m_d(x, t)$ be the number of molecules at time $t$ in the dendritic section $[x, x + dx]$, and let $m_s^{(i)}(t)$ be the number of molecules in the spine connected to the dendrite at $x_i$. We can write this system as

$$\begin{cases} \dfrac{dm_s^{(i)}}{dt}(t) = k_{in}^{(i)} m_d(x_i, t) - k_{out}^{(i)} m_s^{(i)}(t) \\ \dfrac{\partial m_d}{\partial t}(x, t) = D_m \dfrac{\partial^2 m_d}{\partial x^2}(x, t) - \displaystyle\sum_{i=1}^{N} \delta(x - x_i) \dfrac{dm_s^{(i)}}{dt}(t) \end{cases} \tag{S.14}$$

where $D_m$ is the diffusion constant of the considered molecule and $k_{in}^{(i)}$ and $k_{out}^{(i)}$ are the spine-dendrite exchange rate constants pertaining to the $i$-th spine, depending on a number of synaptic and neck features (e.g., width, shape, synaptic crowding and confinement). We associate to this system the no flux boundary conditions $\partial_x m_d(0, t) = \partial_x m_d(L, t) = 0$, as we assume that the influx rate in and outside of the dendrite is low enough compared to the diffusion constant $D_m$. Under these conditions, the system admits the steady-state solution

$$\begin{cases} m_s^{(i)} = \dfrac{k_{in}^{(i)}}{k_{out}^{(i)}} m_d \\ m_d(x_i) = m_d \end{cases} \tag{S.15}$$

with a constant value of $m_d$ throughout the dendrite, and $m_s^{(i)}$ proportional to this value through the ratio of the exchange constants at the $i$-th spine. The solution of our original problem is hereby represented by the histogram of the values $m_s^{(i)}$, corresponding to the distribution density of the ratio $k_{in}^{(i)}/k_{out}^{(i)}$. A crucial step of this derivation is, therefore, the choice of the probability distribution from which the exchange constants are sampled. We examine, in the next paragraphs, the effect of different plausible choices of this distribution.

**Log-normal distribution** A number of different synaptic and neck features have been found to follow a log-normal distribution, so this could represent a first reasonable choice for the statistics of $k_{in}^{(i)}$ and $k_{out}^{(i)}$. From such a scenario, however, it quickly follows that $m_s^{(i)}$ would also follow a log-normal distribution, as the ratio of two log-normally distributed variables is also described by a log-normal distribution. This fact highlights the stability of log-normality in systems with multiplicative noise (as ours), but does not provide insight into how log-normality emerges in the first place from a less specific, possibly additive, noise. We choose, therefore, to move on to more general distribution choices.

**Normal distribution** This distribution represents an appealing choice, due to its ability to naturally emerge in distributions subject to general additive noise. The ratio of normal variables, however, does not have a universally defined, well-behaved distribution and strongly depends on the parameters driving the dividend and the divisor. There is, however, one physical constraint that leads to interesting properties. By definition, $k_{in}^{(i)}$ and $k_{out}^{(i)}$ have to be positive quantities, as they describe positively defined flux rates, into and from the spines. To respect this condition, their distributions will have to have positive means and small enough variances to render the probability of a negative sample negligible. In other words, they have to have a small enough coefficient of variation (CV). Under these conditions, it can be shown that the ratio of two Gaussian variables can be well approximated by a log-normal distribution, with the quality of the approximation decreasing with the magnitude of the coefficients of variation. This approximation, known in the statistical literature as the *delta method*, is valid in general for every function of a narrowly distributed normal random variable. In our case, the derivation starts by considering the variable $Z = \log X/Y$, and approximates it via the Taylor expansion up to the second order

$$Z = \log\left(\frac{X}{Y}\right) \approx \log\left(\frac{\mu_X}{\mu_Y}\right) + \frac{X-\mu_X}{\mu_X} - \frac{Y-\mu_Y}{\mu_Y} - \frac{1}{2}\left[\left(\frac{X-\mu_X}{\mu_X}\right)^2 - \left(\frac{Y-\mu_Y}{\mu_Y}\right)^2\right] \tag{S.16}$$

Remembering that $(X-\mu_X)/\mu_X$ and $(Y-\mu_Y)/\mu_Y$ follow independent normal distributions with mean 0 and variances $CV_X^2$ and $CV_Y^2$ respectively, we can find the first central momenta of $Z$ up to the leading order

$$\begin{aligned}
&\mathbb{E}[Z] \approx \log\left(\frac{\mu_X}{\mu_Y}\right)\\
&\mathbb{E}\left[(Z-\mathbb{E}[Z])^2\right] \approx CV_X^2 + CV_Y^2\\
&\mathbb{E}\left[(Z-\mathbb{E}[Z])^3\right] \approx CV_X^3 + CV_Y^3\\
&\dots
\end{aligned}$$

the general formula for the $n$-th moment being

$$\mathbb{E}\left[(Z-\mathbb{E}[Z])^n\right] \approx \mathbb{E}\left[\left(\tilde{X}-\frac{\tilde{X}^2}{2}+\tilde{Y}-\frac{\tilde{Y}^2}{2}\right)^n\right] \tag{S.17}$$

$$=\sum_{k=0}^{n}\binom{n}{k}\left\{\sum_{j=0}^{k}\binom{k}{j}\left(-\frac{1}{2}\right)^j \mathbb{E}\left[X^{k+j}\right]\sum_{l=0}^{n-k}\binom{n-k}{l}\left(-\frac{1}{2}\right)^l \mathbb{E}\left[Y^{n-k+l}\right]\right\} \tag{S.18}$$

and allowing to show that in general the $n$-th moment is of the order $\mathcal{O}(\max\{CV_X, CV_Y\}^n)$. For small values of $CV_X$ and $CV_Y$ the distribution $Z$ will, therefore, be well approximated by a normal distribution, and the ratio $X/Y$ will consequently be compatible with a log-normal distribution - independently of the mean for $k_{in}^{(i)}$ and $k_{out}^{(i)}$. To give a quantitative evaluation of this compatibility, we resort to computational sampling. For every pair of coefficients of variation $CV_{in}$ and $CV_{out}$ taken in the range $[0.01, 0.22]$, we extract $n_{spines} = 1000$ "synaptic" values of $k_{in} \sim \mathcal{N}(0.5, 0.5\, CV_{in})$ and $k_{out} \sim \mathcal{N}(0.2, 0.2\, CV_{out})$, arbitrarily picking the mean values as they do not impact the approximation. We then test the resulting log-ratio $\log k_{in}/k_{out}$ for normality using the Anderson-Darling test, and reject compatibility for $p < 0.05$. We repeat this procedure $n = 1000$ times and report the final ratio of log-normal compatible samples for each value of $CV_{in}$ and $CV_{out}$ (Fig. S8). As expected, in this range of coefficients of variation, the log-normal compatible fraction results remarkably high (Fig. S8.b,d,c). We do notice, however, that if the two CVs differ by more than 5%, this compatibility drops considerably, with log-ratio distributions acquiring pronounced left or right tails (Fig. S8.a,e). One final source of non-compatibility emerges when the values of the variation coefficients increase beyond $\sim 0.15$: the sampled $k_{in}$ and $k_{out}$ start including negative values, rendering the ratio distribution structurally incompatible with a log-normal distribution (which by definition has positive support). This leads us to conclude that, to model a scenario where the exchange rates are driven by high coefficients of variation, the Gaussian distribution is not a valid modeling choice. To complete our analysis, we therefore switch to another general distribution, the Beta distribution, which is known to be a reasonable approximation of the Gaussian distribution for low CVs, but can also behave as a high CV probability density while maintaining symmetry and a positive, compact support.

**Beta distribution** In order to understand how the (log-)ratio behaves when $k_{in}$ and $k_{out}$ show a higher coefficient of variation, we model their probability density as a Beta distribution

$$\mathrm{Beta}(x;\alpha,\beta) = \frac{x^{\alpha-1}(1-x)^{\beta-1}}{\mathrm{B}(\alpha,\beta)}, \quad x \in [0,1] \tag{S.19}$$

where $\mathrm{B}(\alpha,\beta)$ is the beta function +1

$$\mathrm{B}(\alpha,\beta) = \int_0^1 x^{\alpha-1}(1-x)^{\beta-1}\,dx \tag{S.20}$$

This distribution, due to the compactness of its support, can take arbitrary variation coefficient values, while giving rise to positive and symmetrically distributed values. Using the same procedure illustrated in the previous paragraph, we explore log-normal compatibility of the ratio distribution $k_{in}/k_out$ extending it to a wider range of CV values, confirming that compatibility is reliably retrievable until roughly $CV \sim 0.35$, and decays quickly thereafter due to an increase in the log-ratio curtosis (Fig. S9, in particular panel c). As before, a high degree of log-normal compatibility is found only when $CV_{in}$ and $CV_{out}$ take similar values, with the log-ratio exhibiting left or right skewness otherwise (Fig. S9.a,d).

To confirm the theoretical predictions derived so far, we simulate the system (S.14) as a discrete-time Markov chain. In this framework, the dendrite is subdivided into a finite number of small volumes (of the same size), with or without a connection to a spine. At every time step, each compartment (dendritic volume, synapse) will be able to exchange the resource $m$ with its neighbours, following the transition equations

$$\begin{cases} \Delta m_s^{(i)}(t+1) = \epsilon_{in}^{(i)} m_d(x_i,t) - \epsilon_{out}^{(i)} m_s^{(i)}(t) \\ \Delta m_d(x_j,t+1) = \epsilon_D \left[ m_d(x_{j+1},t) + m_d(x_{j-1},t) - 2m_d(x_j,t) \right] - \delta_{ij} \Delta m_s^{(i)}(t+1) \end{cases} \tag{S.21}$$

where the different $\epsilon$ represent the different transition probabilities deriving from the respective rate constants. The associated Markov transition matrix is

$$\begin{bmatrix} \vdots \\ \Delta m_d(x_{i-1}) \\ \Delta m_d(x_i) \\ \Delta m_s^{(i)} \\ \Delta m_d(x_{j+1}) \\ \vdots \end{bmatrix} = \begin{bmatrix} \ddots & & & & & & & \\ \dots & \epsilon_D & -2\epsilon_D & \epsilon_D & 0 & 0 & 0 & \dots \\ \dots & 0 & \epsilon_D & -2\epsilon_D - \epsilon_{in}^{(i)} & \epsilon_{out}^{(i)} & \epsilon_D & 0 & \dots \\ \dots & 0 & 0 & \epsilon_{in} & -\epsilon_{out}^{(i)} & 0 & 0 & \dots \\ \dots & 0 & 0 & \epsilon_D & 0 & -2\epsilon_D & \epsilon_D & \dots \\ & & & & & & & \ddots \end{bmatrix} \begin{bmatrix} \vdots \\ m_d(x_{i-1}) \\ m_d(x_i) \\ m_s^{(i)} \\ m_d(x_{j+1}) \\ \vdots \end{bmatrix} \tag{S.22}$$

In the simulation, the exchange step is implemented as a multinomial sampling for each compartment, with number of partitions equal to the number of possible transitions. By repeating the same CV sampling procedure carried out for Fig. S8 and S9, we confirm that indeed $m_s^{(i)}$ shows a synaptic distribution compatible with a log-normal density under the hypothesized conditions on the exchange rate coefficients of variation (Fig. S10). Moreover, the simulated process is able to show two additional hallmarks experimentally observed in relation to synaptic size dynamics [14, 15], i.e., the proportionality between the average synaptic molecular content and its average absolute change, as well as the anticorrelation between changes in synaptic molecular amounts between subsequent observation timesteps (Fig. S11).

| Experiment | Parameter | | Factor | Notes |
| --- | --- | --- | --- | --- |
| [7] | Average basally active kinases | $\mu_K$ | 1.2 | In accordance with [16] |
| | Average basally active phosphatases | $\mu_N$ | 0.83 | FK506 hindering effect on CaN binding to substrate [17] |
| | Stimulus induced phosphatases | $N_s$ | 0.55 | Qualitatively following the reasoning for $\mu_N$ |
| [18] | Average basally active kinases | $\mu_K$ | 1.8 | Considering [16] and the higher concentration of FK506 used in experiment compared to [7] |
| | Average basally active phosphatases | $\mu_N$ | 0.53 | FK506 hindering effect on CaN binding to substrate [17], higher FK506 concentration compared to [7] |
| | Stimulus induced phosphatases | $N_s$ | 0.40 | Qualitatively following the reasoning for $\mu_N$ |

Table S3: **Parameter modifications describing FK506 addition.** The modifications are reported for the two works described in the results section. In [7], a concentration of $2\,\mu M$ FK506 was used, while in [18] the concentration amounts to $10\,\mu M$.
