## Supplementary figures for "Push-and-pull protein dynamics leads to log-normal synaptic sizes and probabilistic multi-spine plasticity"

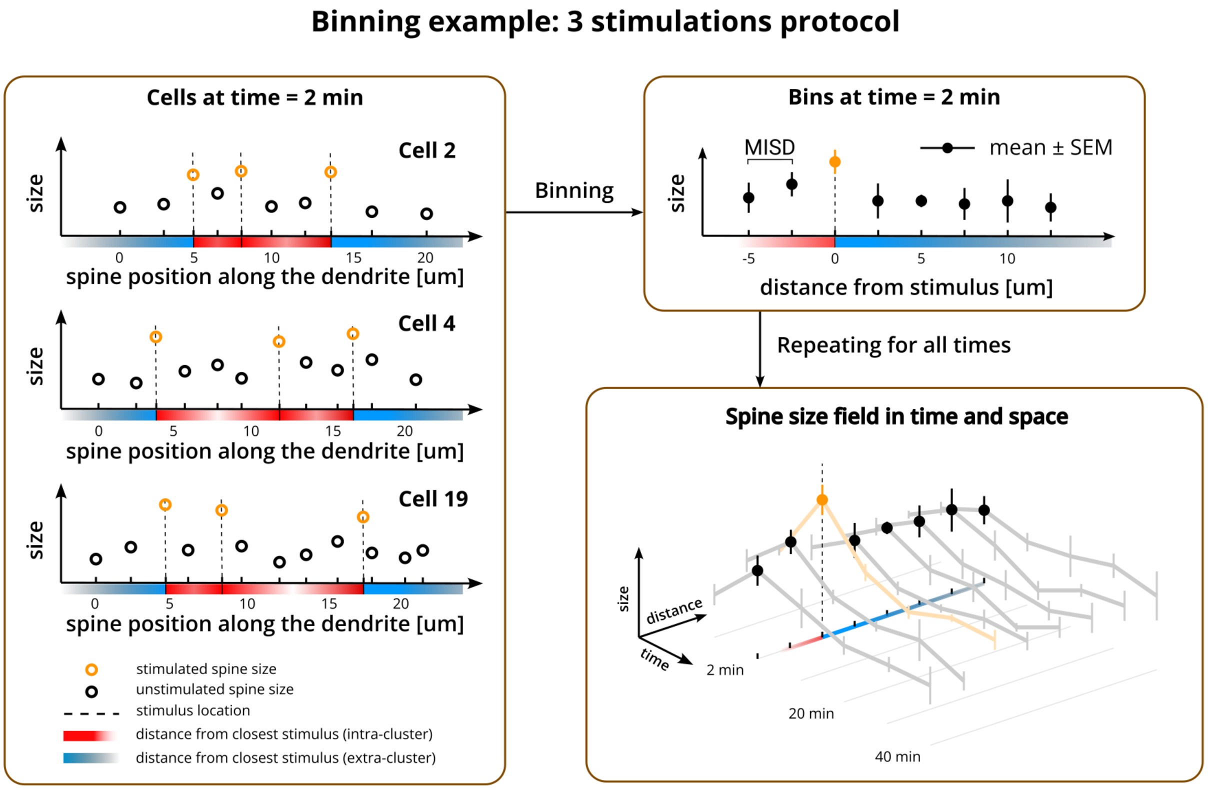


Figure S1: **Graphical representation of the binning procedure.** **a** Considering a specific protocol (e.g., 3 stimulations), for each neuron, at a fixed time point, each spine gets assigned a distance value from the closest stimulus. This value is positive for spines lying outside of the stimulation cluster (blue color code) or negative (red color code). **b** Using bins of width equal to the inferred mean inter-spine (MISD), spine sizes are binned, and the result is reported in terms mean and standard error of the mean. Notice that the resulting "summary" dendrite contains a reduced number of spines. **c** The procedure is repeated for all the considered time points, giving rise to a synaptic size field in function of time and distance from the closest stimulus.

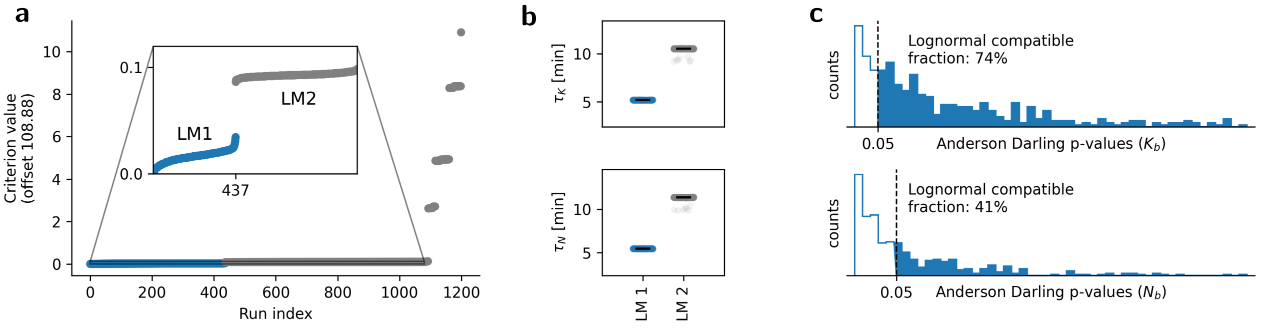


Figure S2: **Model optimization summary.** **a** Cascade plot concerning the 1200 optimization runs. Two (almost equivalent) local minima are found (LM1 and LM2) with convergence basins covering the majority of the available parameter space. **b** Best estimates of the catalytic decay constants $\tau_K$ and $\tau_N$ obtained in the runs converging to the two best local minima. **c** Fraction of compatible $K_b^{(i)}$ and $N_b^{(i)}$ distributions inferred among the runs converging to the first local minimum.

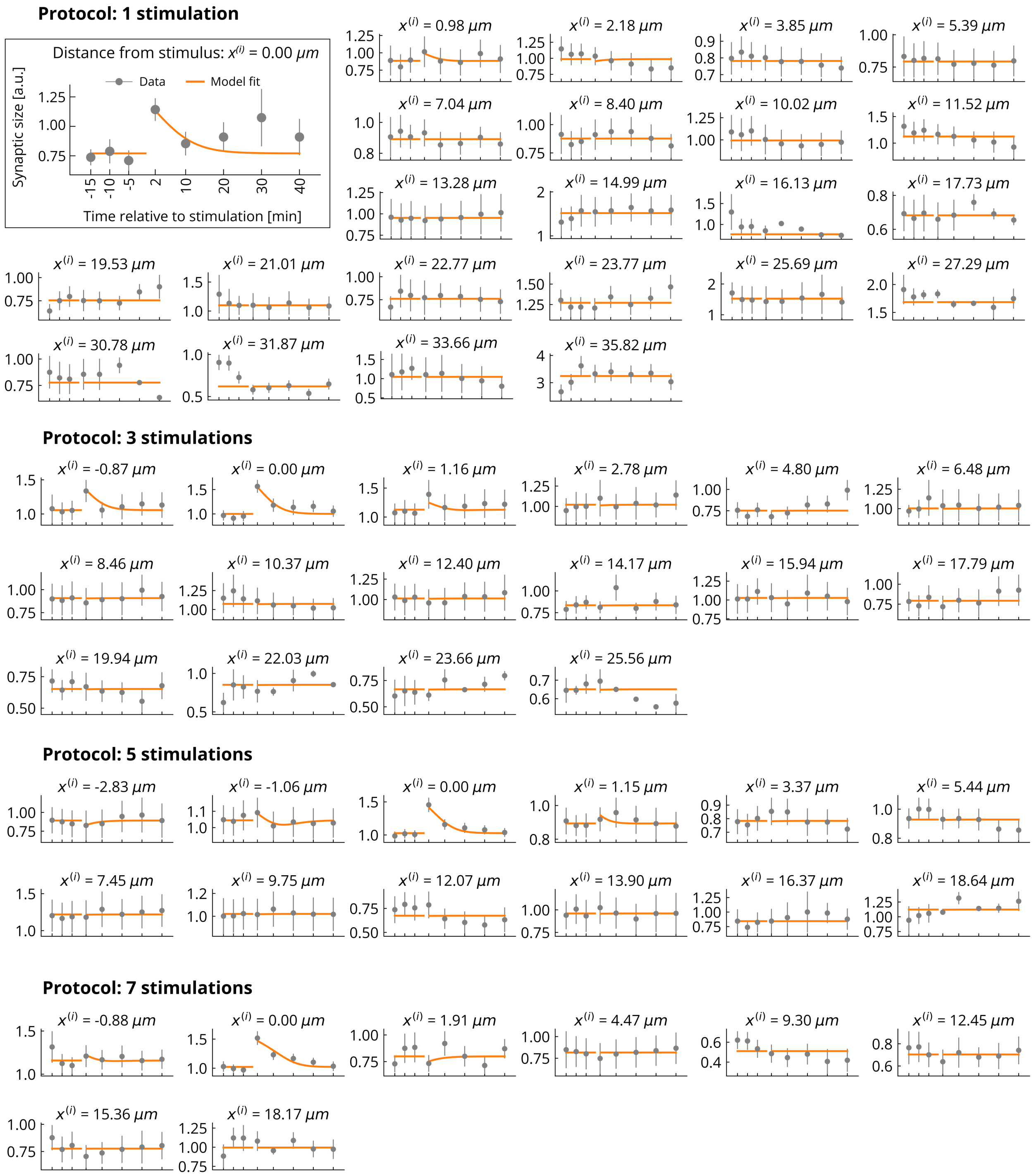


Figure S3: **Fit to data of the optimally parametrized model.** The axis values and the legend are reported only for the bigger, top-left plot, and do not change for the rest of the figure (we removed them for better visualization). The reported values are *absolute* spine sizes (not relative variations) in arbitrary units.

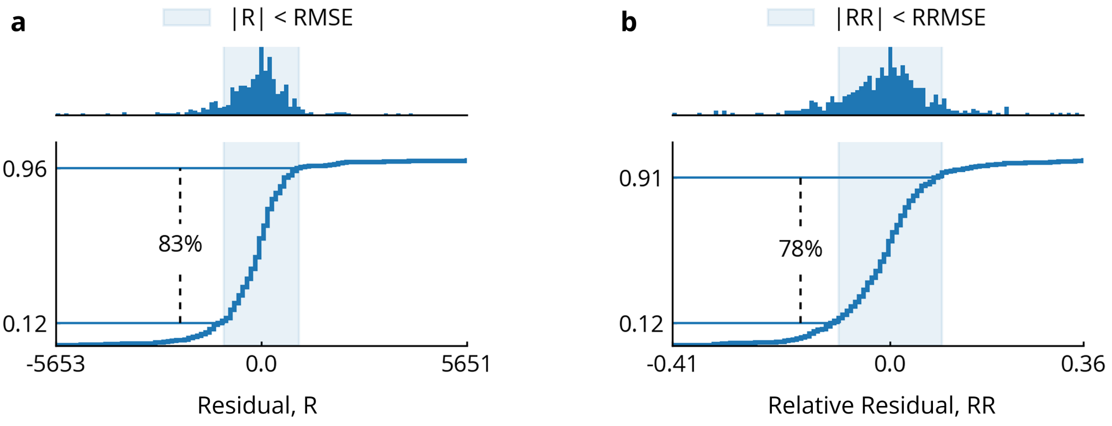


Figure S4: **Optimized model prediction residuals.** **a** Absolute residual (R) probability density (upper graph) and cumulative distribution functions (lower graph) for the best optimization run, 83% of the residuals fall, in absolute value, under the root mean squared error ($RMSE = 1034.51$). **b** Same analysis concerning relative residuals (RR). Importantly, a value smaller than 10% is achieved for the relative root mean squared error ($RRMSE = 9.72\%$), a hallmark of good model convergence.

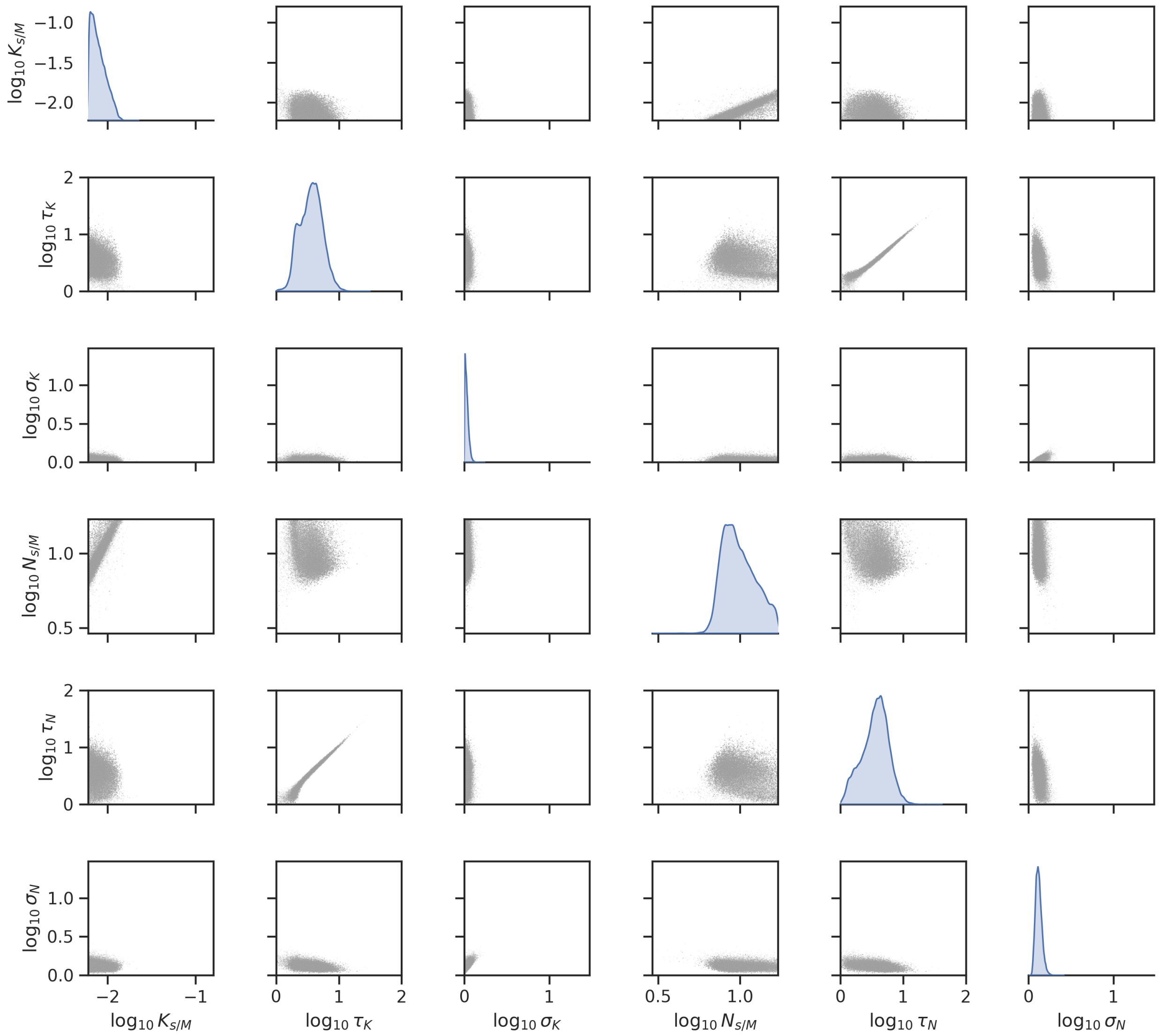


Figure S5: **Posterior distributions of the global dynamic parameters.**

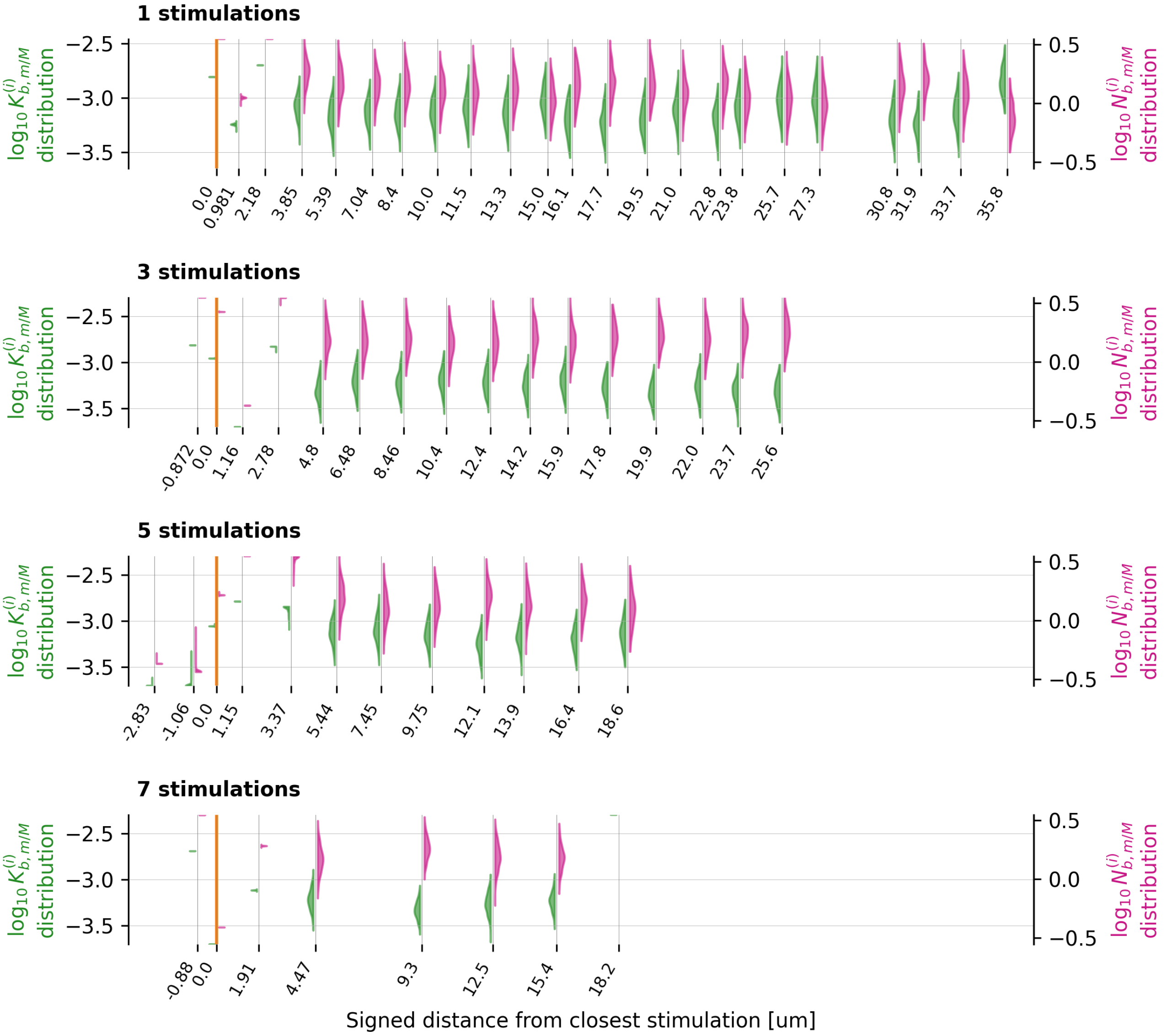


Figure S6: Distribution of inferred optimal values of the basal synaptic parameters across runs of LM1. Each row shows the data obtained from the corresponding protocol. The orange vertical line represents the stimulus location ($0\mu m$ distance).

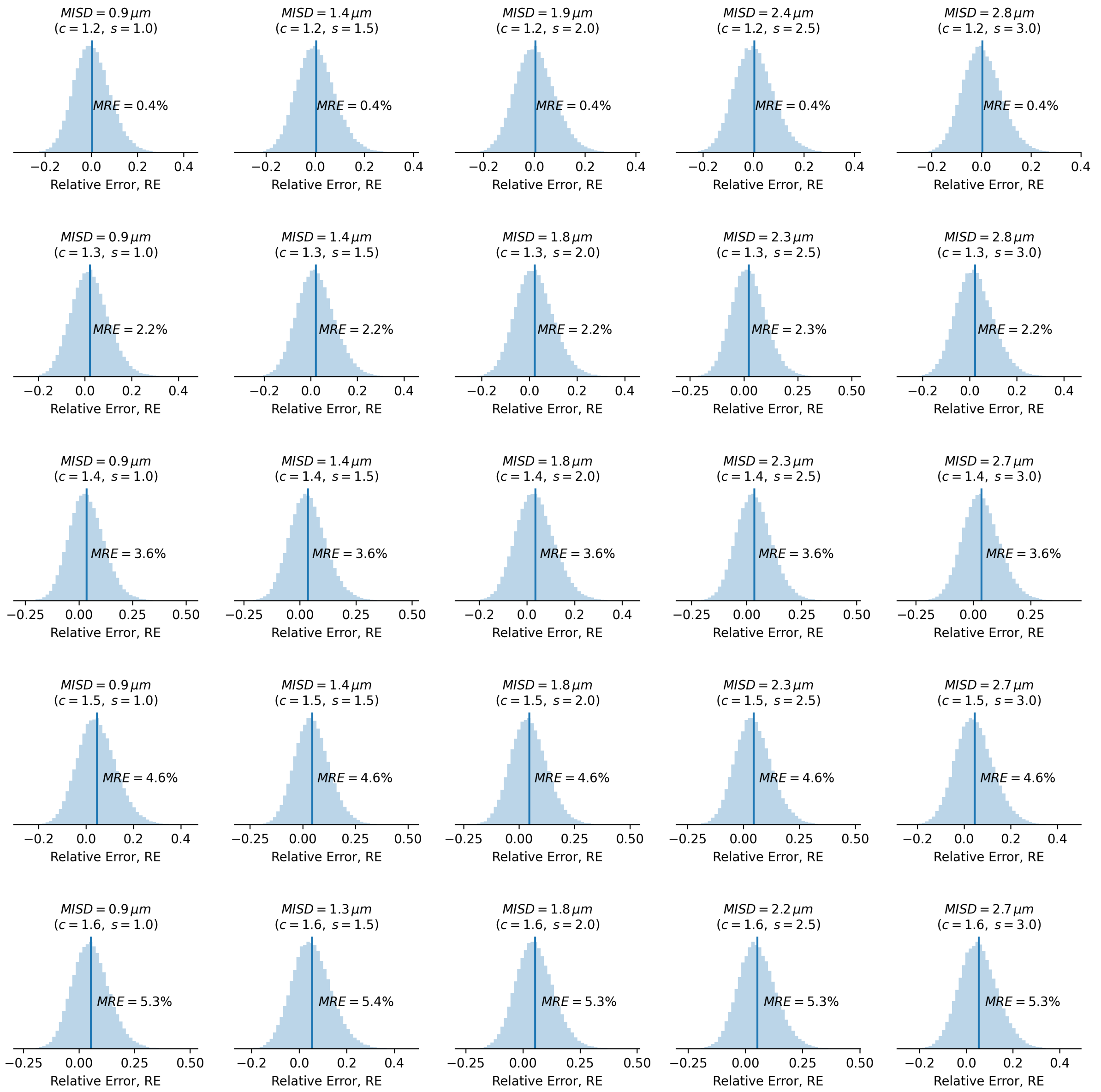


Figure S7: **Relative error sampling of the inter-spine distance Weibull estimator.** The sampling is focused on inter-spine distances compatible with hippocampal spine densities. Notice that the same average inter-spine distance (MISD) can be obtained with different values of the parameters c, s). The vertical bar shows the mean relative error (MRE) of the estimator.

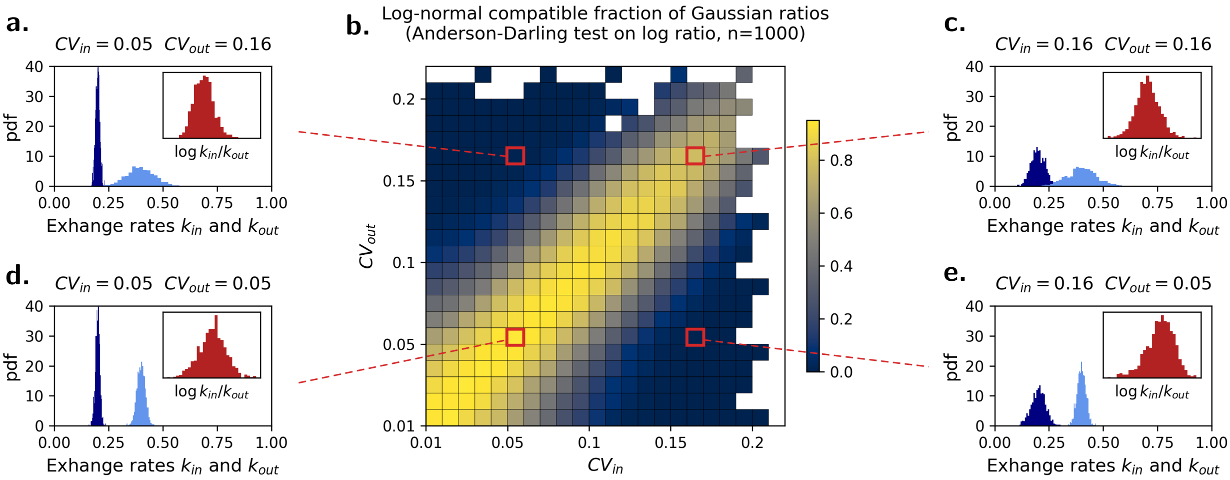


Figure S8: **Log-normal compatibility of ratio distribution generated by normally distributed $k_{in}$ and $k_{out}$.** For each pair of coefficients of variation of $k_{in}$ and $k_{out}$ (referred to as $CV_{in}$ and $CV_{out}$) we sample $n = 1000$ ratios $k_{in}/k_{out}$ and compute the log-normal compatible fraction by testing the log-ratio $\log k_{in}^{(i)}/k_{out}^{(i)}$ for normality with the Anderson-Darling test (panel **b**). As significance threshold, we choose $p = 0.05$. To give a better intuition of the results, we also plot for realizations of different $CV$ values (panels **a,c,d,e**). The while squares in panel **b** represent instances where sampling of $k_in$ and $k_out$ produced values smaller than zero, automatically falsifying log-normal compatibility due to ratios not being positive.

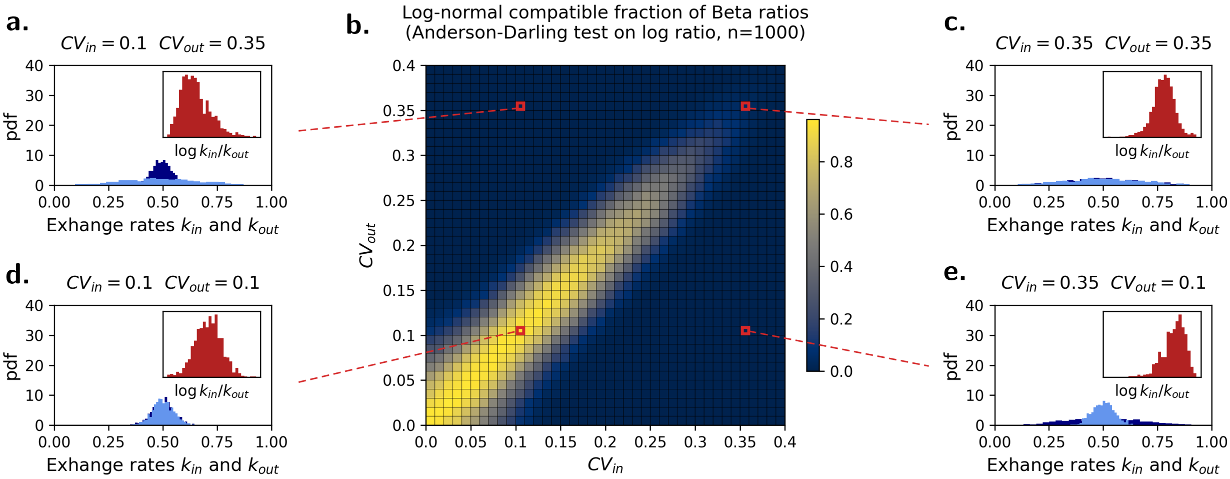


Figure S9: **Log-normal compatibility of ratio distribution generated by Beta distributed $k_{in}$ and $k_{out}$.** The same procedure as in Fig. S8 is carried out. Notice that due to the compact support of the Beta distribution, we are able to explore a wider range of coefficients of variation, finding a considerable degree of log-normal compatibility even in the case of $CV > 30\%$.

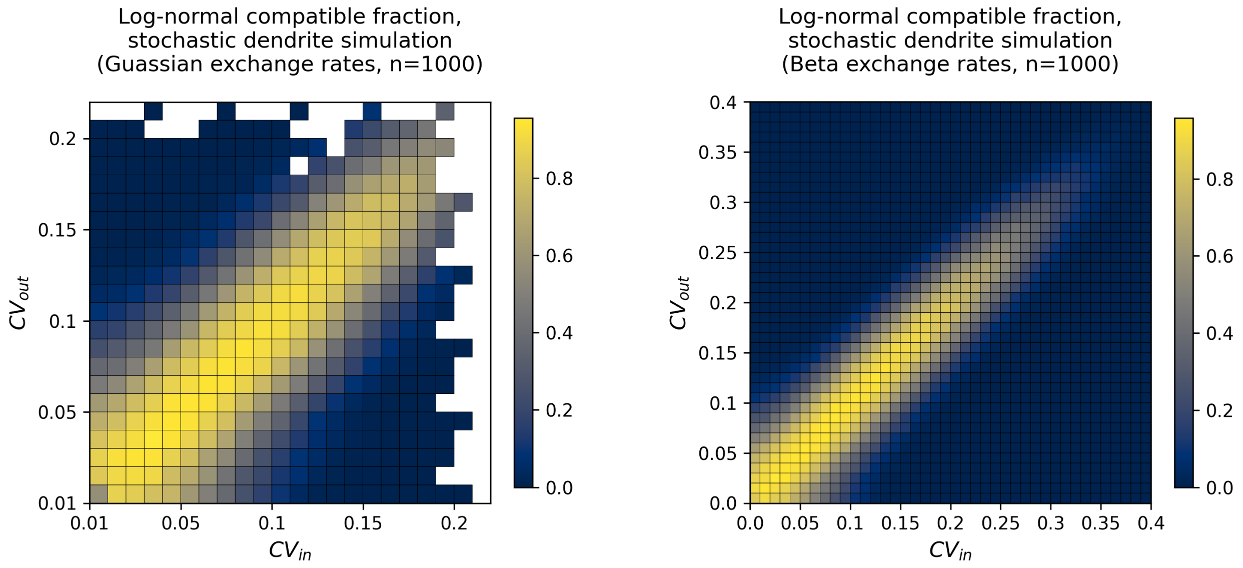


Figure S10: **Log-normal compatibility of stochastically generated synaptic molecular distributions.** The same procedure as in Fig. S8 and S9 is carried out. As hypothesized, stochasticity does not impact the equilibrium synaptic molecular distributions, which follow the theoretically predicted degree of log-normal compatibility.

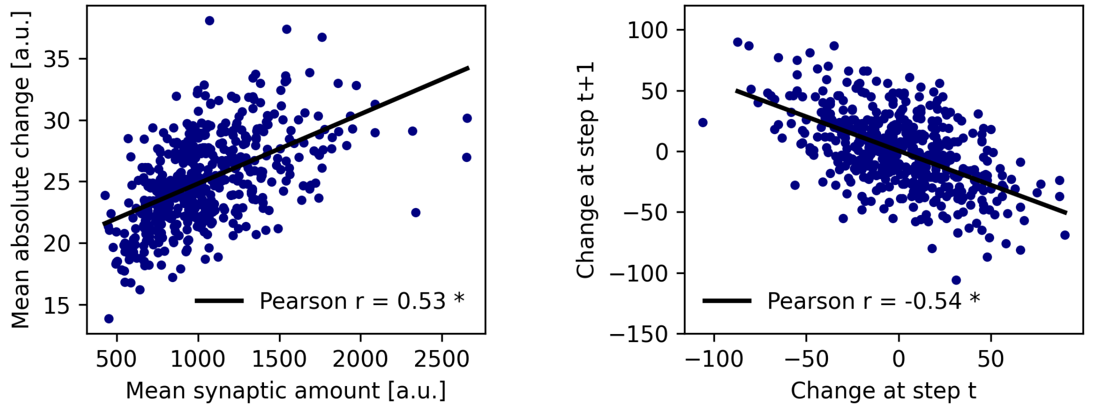


Figure S11: **Analysis of synaptic molecular fluctuations generated with the stochastic simulation.** In accordance with previous reports in the literature, the molecular dynamics produced with the simulations produce two distinctive synaptic features: proportionality between variability and average synaptic amount (left), and anticorrelation of molecular amount changes between subsequent time steps (right).

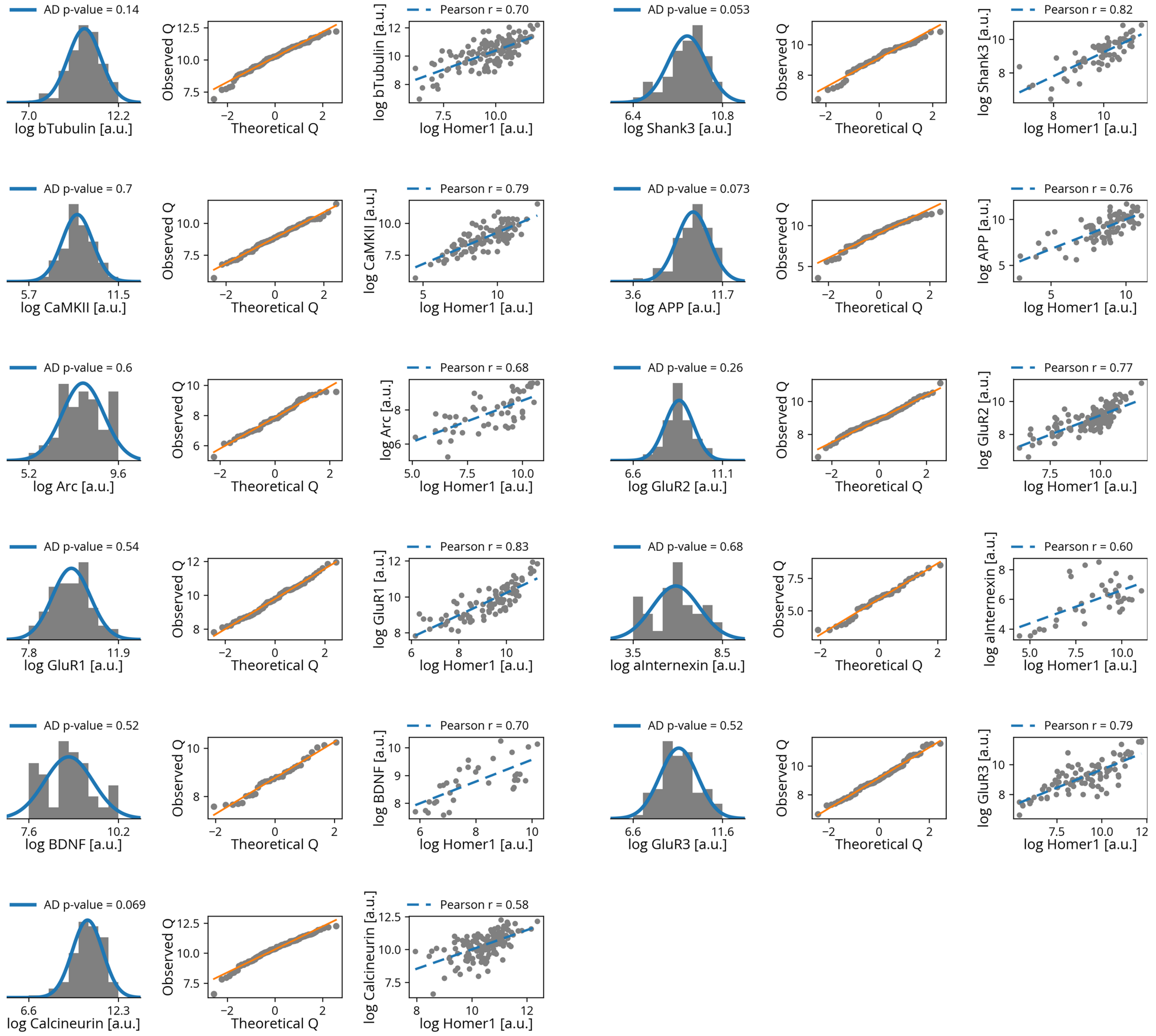


Figure S12: **Distribution, log-normal compatibility and correlation with Homer 1 of several plasticity-related synaptic proteins.**

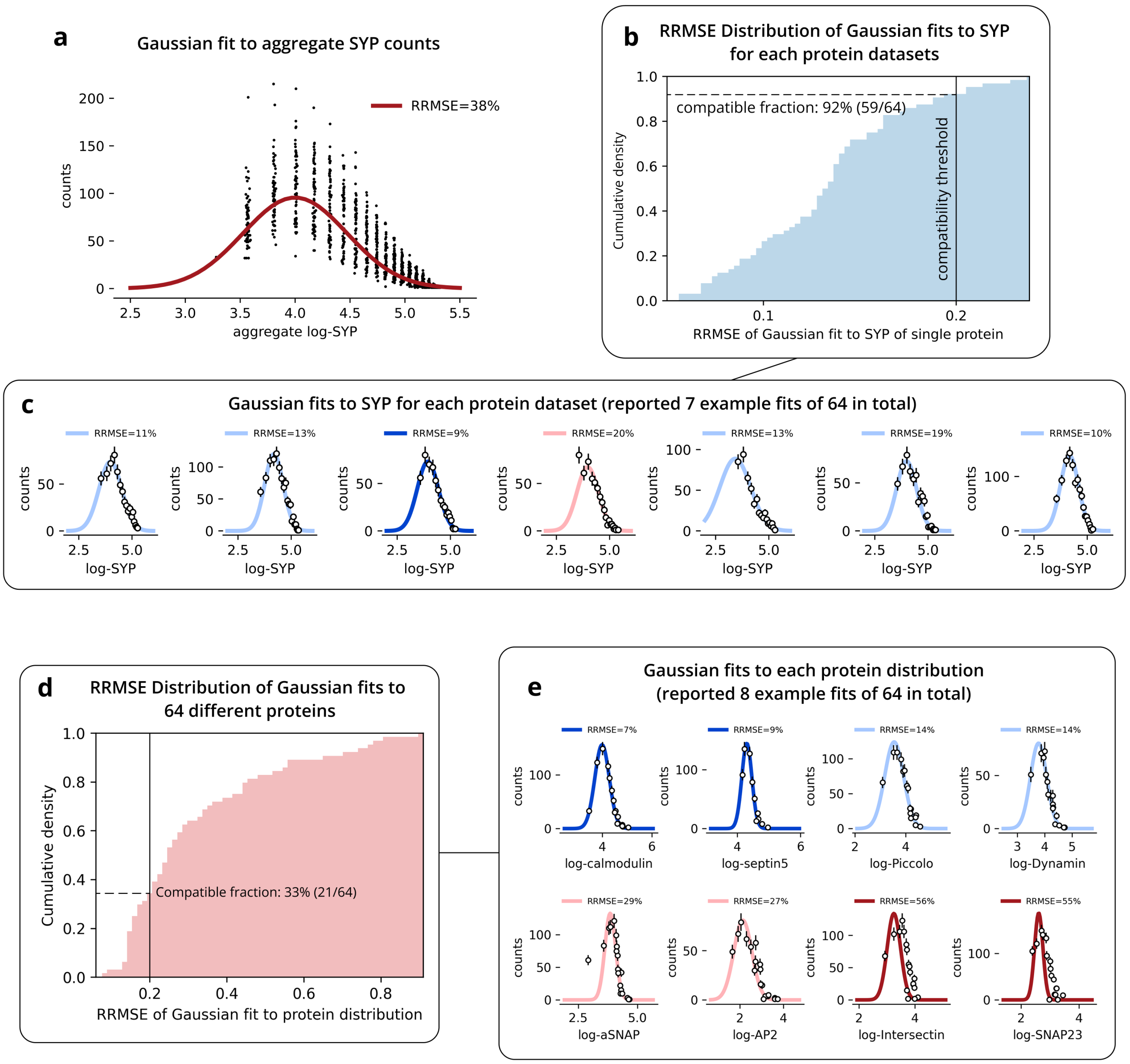


Figure S13: **Log-normal compatibility of pre-synaptic bouton sizes (synaptophysin, SYP) and several pre-synaptic proteins.** RRMSE: relative root mean squared error; compatibility is assessed with RRMSE< 20%. **a** Aggregate SYP counts across different protein datasets. Despite qualitative agreement, log-normal fit to data shows high RRMSE (38%). **b** SYP distributions belonging to each protein dataset show log-normal compatibility for 92% of the datasets. **c** Example fits used to obtain the previous panel. **d** Same as in panel b, but fits carried out on specific protein distributions. In this case, 33% of the proteins result compatible with a log-normal distribution. **e** Example protein distribution fits, showing various degrees of compatibility. Importantly, we can notice that even when the distributions show high RRMSE (pink and red curves), their log-distribution is still qualitatively bell-shaped, suggesting that a higher degree of compatibility could be achieved with different data acquisition.

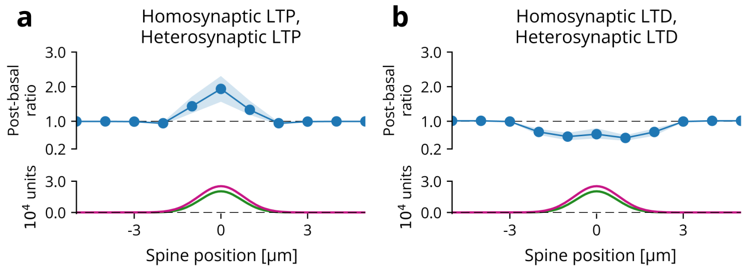


Figure S14: **Different basal catalytic statistics lead to different plasticity response profiles.** For both simulations, we have used the same stimulus parameters leading to Mexican hat potentiation in our main results (the catalytic induction plots are the same for both panels). **a** A reduction of the average basal synaptic content of kinases $\mu_K$ shifts the Mexican-hat potentiation towards a fully potentiating profile. **b** The opposite happens for a reduction in average basal synaptic content of phosphatases $\mu_N$.
